# Exploring the utility of recombinantly expressed snake venom serine protease toxins as immunogens for generating experimental snakebite antivenoms

**DOI:** 10.1101/2022.05.07.491032

**Authors:** Nessrin Alomran, Patricia Blundell, Jaffer Alsolaiss, Edouard Crittenden, Stuart Ainsworth, Charlotte A. Dawson, Rebecca J. Edge, Steven R. Hall, Robert A. Harrison, Mark C. Wilkinson, Stefanie K. Menzies, Nicholas R. Casewell

**Affiliations:** Centre for Snakebite Research & Interventions, Liverpool School of Tropical Medicine, Pembroke Place, Liverpool, L3 5QA, UK; Department of Tropical Disease Biology, Liverpool School of Tropical Medicine, Pembroke Place, Liverpool, L3 5QA, UK; Centre for Drugs and Diagnostics, Liverpool School of Tropical Medicine, Pembroke Place, Liverpool, L3 5QA, UK

## Abstract

Snakebite is a neglected tropical disease that causes high rates of global mortality and morbidity. Although snakebite can cause a variety of pathologies in victims, haemotoxic effects are particularly common and are typically characterised by haemorrhage and/or venom-induced consumption coagulopathy. Despite polyclonal antibody-based antivenoms being the mainstay life-saving therapy for snakebite, they are associated with limited cross-snake species efficacy, as there is often extensive toxin variation between snake venoms, including those used as immunogens for antivenom production. This restricts the therapeutic utility of any antivenom to certain geographical regions. In this study, we explored the feasibility of using recombinantly expressed toxins as immunogens to stimulate focused, pathology-specific, antibodies to broadly counteract specific toxins associated with snakebite envenoming. Three snake venom serine proteases (SVSP) toxins, sourced from geographically diverse and medically important viper snake venoms were successfully expressed in HEK293F mammalian cells and used for murine immunisation. Analyses of the resulting antibody responses revealed that ancrod and RVV-V stimulated the strongest immune responses, and that experimental antivenoms directed against these recombinant SVSP toxins, and a mixture of the three different immunogens, extensively recognised and exhibited immunological binding towards a variety of native snake venoms. While the experimental antivenoms showed some reduction in abnormal clotting parameters stimulated by the toxin immunogens and crude venom, specifically reducing the depletion of fibrinogen levels and prolongation of prothrombin times, fibrinogen degradation experiments revealed they broadly protected against venom- and toxin-induced fibrinogenolytic functional activities. Overall, our findings further strengthen the case for the use of recombinant venom toxins as supplemental immunogens to stimulate focused and desirable antibody responses capable of neutralising venom-induced pathological effects, and therefore potentially circumventing some of the limitations associated with current snakebite therapies.

## Introduction

Snakebite is a significant public health issue, as more than 5.4 million people are bitten annually, resulting in as many as 1.8 million envenoming and 138,000 deaths, which primarily affect impoverished communities in the rural tropics and subtropics across Africa, the Middle East, Americas, Asia and Australasia (1-3). Since 2017, snakebite has been classified by the World Health Organization (WHO) as a neglected tropical disease (4), and in 2018 the WHO outlined a global roadmap with the goal of halving snakebite mortality by 2030 (5).

Because the toxin composition of venomous snakes varies extensively among species (6, 7), the pathological patterns of envenoming also vary, though these can be broadly classified into three major categories: neurotoxic, cytotoxic and haemotoxic (2, 8). Haemotoxicity is one of the most common critical signs observed in envenomed snakebite victims and is particularly common following bites by viperid snakes. Haemotoxic envenoming can result in local and/or systemic haemorrhage, including overt bleeding, such as from the gums or the bite site, and internal bleeding, such as intracranially (2, 8). Such envenoming can also cause coagulopathy, defined ultimately by defibrinogenation, and known as venom-induced consumption coagulopathy (VICC). Coagulopathy can contribute extensively to the severity of envenoming by rendering victims particularly vulnerable to haemorrhage (2, 8-10).

Snakebite coagulopathy is the consequence of certain venom toxins consuming and/or increasing the abnormal activation of key clotting factors (e.g. Factor V, X, II [prothrombin]), ultimately leading to a loss of clotting capability (11-13). Although *in vitro* such ‘procoagulant’ venom toxins cause rapid clot formation, their cumulative effects lead to severe and rapid factor consumption *in vivo*, characterised by depletion of fibrinogen and accompanied by a consequent increased risk of bleeding in victims (10, 14). Many potent procoagulant toxins act by directly activating Factor V, X or II, though others act directly on fibrinogen in a fibrinogenolytic manner (11), and thus contribute towards the consumption of fibrinogen (hypofibrinogenaemia). Many such venom toxins are known as thrombin-like enzymes (TLEs), because these proteins are capable of cleaving the *α*-chain and/or *β*-chain of fibrinogen in a manner analogous to thrombin (15). However, cleavage by TLEs does not result in the liberation of active fibrin (unlike that caused by thrombin), and thus serves to deplete circulating fibrinogen (15). The TLEs are members of the snake venom serine protease (SVSP) gene family, an important toxin class, particularly in viperid venoms (16). This multi-locus gene family typically encodes multiple related isoforms in the venom gland of each snake species, and these toxins can exhibit distinct functional activities other than those associated with fibrinogenolysis (6, 8).

Serum-derived, polyclonal antibody-based antivenoms remain the gold standard therapeutic option for treating snakebite envenoming (17). Despite antivenoms being life-saving therapeutics, they are also associated with several limitations restricting their utility. First, antivenom antibodies are derived from venom-immunised animals, and thus come with the risk of stimulating adverse reactions, which can range from vomiting, urticaria and/or generalised rash to anaphylaxis (18-21). Antivenoms also typically exhibit limited cross-snake species efficacy, as the direct result of variation in venom composition among medically important snake species (6, 22). Thus, antivenoms are typically only effective against snake species whose venoms are highly similar to those included in the immunising mixture. In addition, one major problem with the majority of current antivenoms is that only a relatively small proportion of the active components are typically specific to the venom immunogens (i.e. 10-20% of the IgG or IgG-fragment antibodies) (23). In terms of implementation, antivenoms (i) have to be delivered intravenously and thus require a clinical environment for patient care (also due to the management of potential adverse reactions), and (ii) are often unaffordable or unavailable to the patient in low or middle-income countries, where treatment courses have been reported to cost between $60-$640 USD (24), an expense that can push many victims further below the poverty line (25, 26).

As a result of the aforementioned limitations, several attempts have been made to improve the cross-reactivity and dose efficacy of snakebite treatments over recent years, both via the optimisation of conventional polyclonal antivenoms (e.g. optimisation of venom immunogen mixtures, use of recombinant proteins or epitopes as immunogens) (25-27) and the exploration of new treatment formats, including monoclonal and oligoclonal antibodies and small molecule drugs (28, 29). While the latter approaches show great promise for the long-term future of snakebite therapy, there is a pressing need to improve the efficacy of conventional therapy in the short term. This study explores the potential tractability of using recombinantly expressed snake venom proteins, specifically SVSP toxins, as immunogens for generating focused toxin-specific antibody responses.

The expression of recombinant proteins has been historically undertaken in a wide variety of different host systems, including bacteria, yeast, plant, insect and mammalian cells. Several snake venom proteins, including ancrod from *Calloselasma rhodostoma* venom (30), acutin from *Deinagkistrodon acutus* (31), batroxobin from *Bothrops atrox* (32), and factor V activator from *Macrovipera lebetina* (33) have previously been expressed in either bacteria (e.g. *E. coli*) or yeast (e.g. *Pichia pastoris*). However, some snake venom toxins can be challenging to successfully express in such systems because of the frequency of cysteine residues and disulfide bonds, resulting in significant obstacles relating to correct protein folding, solubility and yield (23, 34), even under conditions intended to favour disulfide bond formation (35, 36). Further, *E. coli* lack glycosylation apparatus limiting their use if effector functions are needed (37), while expression in yeast may result in different glycosylation patterns in comparison with the native protein (30). Consequently, over the past decade, the use of transient mammalian expression systems has increased, and have been applied to produce recombinant snake venom toxins, such as the SVSP gyroxin (found in native form in *Crotalus durissus terrificus* venom) (38). Mammalian expression systems offer a number of desirable characteristics for the expression of vertebrate venom toxins (i.e. native protein folding and post-translational modifications) (39-41), and Human Embryonic Kidney 293F (HEK293F) cells in particular offer ease of transfection, high expression yields and native human glycosylation amenable for such work.

In this study, three fibrinogenolytic SVSPs sourced from distinct medically important viperid snake venoms (ancrod from *Calloselasma rhodostoma*, batroxobin from *Bothrops atrox* and RVV-V from *Daboia russelii*) were expressed in HEK293F mammalian cells for use as immunogens to generate experimental antivenoms. To this end, following functional validation of the resulting recombinantly expressed toxins in fibrinogenolysis experiments, a murine immunisation regimen was undertaken to generate polyclonal antibodies against each of these three toxins. Thereafter, we explored the *in vitro* immunological recognition and inhibitory capability of the resulting antibodies against the different recombinant toxin immunogens and a panel of crude snake venoms. Our findings demonstrated that polyclonal antibodies generated against specific venom toxins exhibited broad *in vitro* cross-reactivity with a geographically diverse array of native snake venoms and were capable of inhibiting certain toxin functional activities, including fibrinogenolysis and the prolongation of plasma clotting times. More broadly, our findings suggest that the recombinant expression of key, functionally important, snake venom toxins, could be a valuable approach to generate new pathology-specific antivenoms against snakebite or to enhance existing antivenoms via increasing antibody titres against specific pathogenic toxins.

## Material and methods

### Selection of SVSPs toxins for expression

Three SVSP toxins were selected for recombinant expression due to their prior characterisation as biologically active and functionally relevant components of distinct viper venoms, specifically: ancrod from *C. rhodostoma* (Genbank: L07308.1) (42), batroxobin from *B. atrox* (Genbank: J02684.1) (43) and RVV-V from *D. russelii* (Genbank: MF289120.1) (34). Coding sequences were sourced from the GenBank database of the National Centre for Biotechnology Information (http://www.ncbi.nlm.nih.gov/genbank/), and the SignalP-5.0 Server (http://www.cbs.dtu.dk/services/SignalP/website) was used to detect and remove the presence of signal peptides. Integrated DNA Technologies (IDT, UK) commercially synthesised sequences of inserts (ancrod, batroxobin and RVV-V) for cloning into the expression vector (Supplementary Figure 1).

### Vector and insert restriction digests

For the generation of plasmid vectors containing the toxin-encoding DNA inserts, we used commercially available pFUSE-hlgG1-Fc2 Plasmids (4194 bp) (Invitrogen, USA). Restriction digests were performed using 5 µg of the pFUSE-hlgG1-Fc2 vector, 10 µl of 10X buffer 2.1 (New England BioLabs®Inc., UK), 5 µl of each restriction enzyme EcoRI (20,000 U/ml; cut site: 5’-GAATTC-3’) and NheI (10,000 U/ml; cut site: 5’-GCTAGC-3’) (New England BioLabs®Inc., UK) and 100 µl of nuclease-free water, followed by incubation for one hour at 37 °C in a water bath. Following brief centrifugation, dephosphorylation of the resulting digested vector was performed by the addition of 20 µl of shrimp alkaline phosphatase (1,000 U/ml, New England BioLabs®Inc., UK), 20 µl of 10X CutSmart buffer (New England BioLabs®Inc., UK) and 60 ml nuclease-free water, followed by a further one-hour incubation step at 37°C. Thereafter, samples were briefly centrifuged again and incubated for 30 minutes at 65 °C to deactivate the alkaline phosphatase. DNA inserts (ancrod, batroxobin and RVV-V) (Integrated DNA Technologies, Inc. (IDT), UK) were prepared using 500 ng of insert, 10 µl of 10 x buffer 2.1: (50 mM NaCl, 10 mM Tris-HCl, 10 mM MgCl_2_ and 100 µg/ml BSA pH 7.9), 5 µl of the EcoRI (20,000 U/ml) and NheI (10, 000 U/ml) restriction enzymes, and 100 ml of nuclease-free water, followed by incubation for one hour at 37 °C in a water bath and brief centrifugation thereafter.

Digests for both vector and inserts were examined on 0.8% agarose (Sigma-Aldrich, UK) in 1x TBE buffer (0.089 M Tris-Borate, 0.002 M Ethylenediaminetetraacetic acid (EDTA), pH 8.3). The solution was heated in a microwave until fully dissolved before the addition of 9 µl of ethidium bromide (Sigma-Aldrich, UK) and polymerisation in a Bio-Rad gel system. Gels were immersed in 1x TBE buffer prior to the loading of 50 μl and 200 μl of vector and insert, respectively, alongside 10 μl (0.5 μg) of 1 kb DNA ladder (New England BioLabs®Inc., UK). The running conditions were 50 V, 240 mA for 80 minutes, with downstream visualisation using a Gel Doc EZ Gel Documentation System (Bio-Rad).

### Plasmid DNA Purification

For DNA purification, the bands representing each insert and the vector were dissected from the gel and placed in 15 ml Falcon™ Conical Centrifuge Tubes, where they were weighed, and 3 x volumes of solubilization buffer QG (QIAquick PCR Purification Kit, QIAGEN, Germany) added to 1 x volume of gel (100 mg, ∼100 µl), before incubation for 5-10 minutes at 56 °C in a water bath. Next, 2 x volumes of isopropanol to 1 x volume of gel was added to each sample before the solution was transferred to a QIAprep 2.0 Spin Miniprep column (QIAquick PCR Purification Kit, QIAGEN, Germany), centrifuged briefly at 2817 x g, and the eluate discarded. Next, 500 µl of QG buffer was added to each column, followed by centrifugation at 2817 x g for 1 minute, and then 700 µl of PE buffer (The washing buffer, QIAquick PCR Purification Kit, QIAGEN, Germany) was added followed by centrifugation at 2817 x g for 1 minute, with the eluate discarded after each step. Following a final dry spin at 2817 x g for a few seconds, each column was transferred into an Eppendorf tube before 50 µl of elution buffer (QIAquick PCR Purification Kit, QIAGEN, Germany) was added, samples incubated at room temperature for two minutes, before centrifugation for 1 minute at 2817 x g. The resulting insert and vector DNA concentrations were quantified by Nanodrop and the samples stored at -20 °C until downstream use.

### Plasmid ligation, amplification and purification

The inserts were ligated into the vector using a 3:1 molar ratio. The necessary volumes of each insert for ligation with 1 µl of vector were calculated (http://www.insilico.uni-duesseldorf.de/Lig_Input.html) and prepared in ligation reactions consisting of 1 µl of 10x ligase buffer (supplied in 300 mM, Tris-HCl (pH 7.8), 100 mM MgCl2, 100 mM DTT and 10 mM ATP) and 1 µl of T4 ligase (supplied in 10 mM Tris-HCl (pH 7.4), 50 mM KCl, 1 mM DTT, 0.1 mM EDTA and 50% glycerol) (both Promega, UK), with the final reaction volume adjusted to 10 µl with nuclease-free water. Control reactions were also performed in parallel and consisted of vector alone without the insert. Ligation reactions were incubated at 16 °C overnight. Next, ligated DNA was transformed using TOP10 *E. coli* competent cells (Thermo-Fisher Scientific, UK). Competent cells were thawed on ice for 10 minutes, 50 µl of cells added to 5 µl of each ligation mixture and incubated on ice for 30 minutes, followed by heat shock for 30 seconds at 42 °C in a water bath. The bacteria were then incubated on ice for two minutes before adding 250 µl pre-warmed SOC Outgrowth Medium (New England BioLabs®Inc., UK) and incubating in a shaking incubator at 220 rpm, 37 °C, for 1 hour. Thereafter, bacteria were streaked onto LB plates (6 g LB Broth (Luria low salt) (Sigma-Aldrich, UK), 4.5 g agar (Sigma-Aldrich, UK), 300 ml water, pH 7.5-7.8, containing 75 µl Zeocin (25 µg/µl, InvivoGen, UK)) before incubation at 37 °C overnight. A single bacterial colony for each sample was then selected and used to inoculate 5 ml of LB media, before shaking incubation at 37 °C for 6-8 hours at 220 rpm. Erlenmeyer flasks (125 ml) were prepared with 50 ml of LB media supplemented with 12.5 µl Zeocin (25 µg/µl, InvivoGen, UK). A total of 1 ml of resulting bacteria was added to each flask before shaking incubation at 37 °C and 220 rpm overnight. Thereafter, glycerol stocks were prepared using a 300 µl aliquot of the overnight bacterial culture mixed with 700 µl sterile 50% glycerol before storage at -80 °C.

Plasmid DNA was purified using the Hi-Speed Plasmid Midi Kit (QIAGEN, Germany). Overnight cultures were centrifuged at 6000 x g for 10 minutes at room temperature. Pellets were then resuspended in 6 ml of Buffer P1 (50 mM Tris-HCl pH 8.0, 10 mM EDTA, 100 μg/ml RNaseA) before 6 ml of Buffer P2 (lysis buffer: 200 mM NaOH, 1%) was added, and the solution mixed thoroughly by inverting gently and standing for five minutes. Next, 6 ml of Buffer P3 (neutralisation buffer: 3.0 M potassium acetate (KAC), pH 5.5) was added and mixed thoroughly by gently inverting it several times. The lysate was added to a QIAfilter Cartridge and incubated for 10 minutes. Meanwhile, 4 ml of Buffer QBT (equilibrium buffer: 750 mM NaCl, 50 mM 3-(N-morpholino) propane sulfonic acid (MOPS) pH 7.0, 15% isopropanol, 0.15% triton X-100) was added to a HiSpeed Tip and incubated for 10 minutes. A plunger was inserted into the cartridge, and the solution with the plasmid DNA was transferred to the HiSpeed Tip. Retained plasmid DNA was washed with 20 ml of Buffer QC (1.0 M NaCl, 50 mM MOPS, pH 7.0, 15% isopropanol) and eluted with 5 ml of Buffer QF (1.25 M NaCl, 50 mM Tris-HCl pH 8.5, 15% isopropanol). For DNA precipitation, 3.5 ml of isopropanol was added (final concentration 70%), mixed, and incubated for five minutes. Next, the precipitated plasmid DNA solution was expelled through a QIprecipitator using constant pressure and then washed with 2 ml 70% ethanol before air drying. Finally, 600 µl of elution buffer (buffer EB, 10 mM Tris-Cl, pH 8.5) was added to the QIprecipitator to elute the DNA. To determine the resulting DNA yield, DNA concentrations were measured via Nanodrop (Thermo-Fisher Scientific, UK). DNA plasmids were then commercially sequenced via Sanger sequencing (Source Bioscience, UK) to confirm insert fidelity.

### Expression in HEK293F cells

For protein expression, we used HEK293F cells and the Freestyle 293 Expression System (Thermo-Fisher Scientific, UK). Cells were maintained in suspension, in Freestyle serum-free media, with culture conditions of 37 °C, 125 rpm and 8% CO_2_. Cells were originally seeded at 2 × 10^5^/ml and grown over three days to 1 – 2 × 10^6^/ml. Cell cultures were counted and checked for at least 90% viability using a Luna II automated cell counter (Labtech, UK). The cells were seeded at 7 × 10^5^/ml for transfection, equivalent to 2.1 × 10^7^ cells total in 30 ml medium. Cells were left to grow for 16 – 24 hours. 30 mg of DNA was placed into a sterile Eppendorf tube for each Erlenmeyer flask (125 ml). An equivalent volume of Freemax transfection reagent was added to a separate tube, and the volumes of each made up to 600 ml with Optipro SFM transfection medium (Gibco). Both 600 ml aliquots were mixed together and incubated for 12 minutes at room temperature. The mixture was then added to swirling cells in the Erlenmeyer flasks (125 ml), and the cells were left to grow for seven days at 37°C, 8% CO_2_ on a shaking platform. After seven days, cell cultures were pooled and centrifuged at 500 × g for five minutes to pellet the cellular debris. Supernatants were filtered through a 0.22 µm pore membrane and stored at 4 °C before protein purification. Proteins were purified by hydrophobic affinity chromatography using an ÄKTA pure protein purification system (GE Healthcare). Lysates were loaded onto a 1 ml HiTrap™ Chelating Sepharose™ Nickel column (GE Healthcare) in 40 mM imidazole in Na phosphate buffer 20mM with 150mM NaCl pH 7.4 and eluted with 500 mM imidazole, 150 mM NaCl, 20mM sodium phosphate buffer pH 7.4. The fractions were pooled and dialysed in Slide-A-Lyzer G2 Dialysis Cassettes (1-30 ml) 2 K molecular weight cut-offs (MWCOs) (Thermo-Fisher Scientific, UK). This step was performed via three changes of 1x PBS (pH 7), first for two hours at room temperature, secondarily at room temperature overnight and then for two hours at room temperature again. Finally, the protein was concentrated using 30 K MWCOs concentrator tube (Thermo-Fisher Scientific, UK) and centrifuged at 1684 x g for 15-30 minutes. Protein concentrations were quantified by Bradford protein assay (Bio-Rad, UK) by using 96-well flat-bottomed plates (NUNC) and a LabTech LT-4500 microplate absorbance reader at 595nm.

### Murine immunisations

Next, we used each recombinantly expressed snake venom toxin in immunisation experiments. Prior to immunisation, we performed quality control experiments on the recombinant toxins, namely SDS-PAGE gel electrophoretic assessments of purity, and fibrinogenolytic and clotting time assessments of functional activity, using the methods outlined in full below. The murine immunisation experiments were undertaken with the approval of The Animal Welfare and Ethical Review Boards of the Liverpool School of Tropical Medicine and the University of Liverpool, under project license (#P58464F90) granted from the UK Home Office, and in accordance with the UK Animals (Scientific Procedures) Act 1986.

Murine immunisation experiments broadly followed a recently outlined approach (44). Mixed-gender experimenters performed all experiments. Experimental animals were housed in Techniplast GM500 cages with Lignocel bedding (JRS, Germany) and zigzag fibres nesting material (Sizzlenest, RAJA), and kept at approximately 22 °C at 40-50 % humidity, with 12/12 hour light cycles. Experimental animals were kept in specific-pathogen-free facilities with ad libitum access to certified reference materials, including irradiated food (Special Diet Services) and reverse osmosis water (in an automatic water system). All cages were changed every two weeks. Animals were monitored twice per week throughout the course of immunisation for signs of adverse reactions (e.g., reduced activity, physiological impairment, pallor, ulceration following immunisation) and general health (e.g., loss of body weight), and no animals were culled due to weight loss or behavioural endpoints being met.

Experimental animals were allowed to acclimatise for one week, and thereafter groups of five female CD-1 mice (initial weight 25-30 g; Charles River, UK; randomised to groups) were immunised with either: (i) recombinant ancrod, (ii), recombinant batroxobin, (iii) recombinant RVV-V, (iv) a 1:1:1 mixture of the three recombinant toxins, or (v) a positive control group consisting of a 1:1:1 mixture of three crude snake venoms sourced from *Calloselasma rhodostoma* (captive bred), *Daboia russelii* (Sri Lanka) and *Bothrops atrox* (Brazil). Lyophilised snake venoms were sourced from the historical venom collection stored in the herpetarium of the Centre for Snakebite Research and Interventions (CSRI) at the Liverpool School of Tropical Medicine. All experimental animals received 1 µg of toxin/venom at each immunisation (except for the batroxobin group, which received 2 µg for the third and fourth immunisation followed interim evaluation of serological responses), with the groups receiving the mixture of recombinant toxins or venoms receiving equal amounts (1:1:1 ratio, i.e., 0.33 µg per toxin/venom, 1µg total) of the different toxins/venoms per contributing sample. All immunogens were prepared in sterile PBS and mixed in a 1:1 ratio with the Sigma Adjuvant System (Sigma-Aldrich, UK), to a final injection volume of 40 µl.

Each experimental animal was immunised a total of five times, with secondary immunisations occurring two, four, eight and twelve weeks after primary immunisation. For immunisation, animals were briefly anaesthetised with an inhalational anaesthetic (isoflurane, 5% for induction of anaesthesia and 1.5 to 3% isoflurane for maintenance of anaesthesia). Whilst under anaesthesia, the dorsal fur was shaved at the injection site, and all animal groups were subcutaneously injected with 1 µg immunogen in a total volume of 40 µl, with the first two immunisations delivered to a single injection site and thereafter split across two sites (i.e., 20 µl/site). Interim blood samples to profile immune responses were collected via tail tip sampling on weeks three, six, ten and a terminal sample was collected following euthanasia at week 14. All blood samples were allowed to clot for a minimum of two hours at room temperature and centrifuged at 23548 x g for 10 minutes. The resulting sera were stored at -20 °C until further use.

### IgG Purification

The terminal murine serum obtained for each group were pooled and then IgG purified to produce the resulting experimental antivenoms (anti-ancrod, anti-batroxobin, anti-RVV-V, anti-toxin mix, anti-venom mix). Pre-immunisation serum samples were processed in an identical manner to serve as the negative control (i.e., non-immunised). IgG was purified from pooled mice serum (27) using 14 week (terminal) blood samples, which were first diluted with 0.90% saline (1:1 ratio), before the slow (drop by drop) addition of caprylic acid (Octanoic acid; Sigma-Aldrich, UK) with stirring to a final concentration of 6%. Subsequently, non-immunoglobulin proteins were precipitated by vigorous stirring for 60 minutes, followed by centrifugation at 12800 x g for 60 minutes at 4°C. The resulting supernatant from each sample was dialysed separately using 35 mm 3.5K MWCO SnakeSkin Dialysis Tubing (Thermo-Fisher Scientific, UK) and sodium citrate buffered saline (SCS, pH 6.0), with two changes of SCS, first at room temperature for one hour, and then secondarily at 4 °C overnight. Following dialysis, the final IgG samples were lyophilised using a Labogene lyophiliser and stored at 4°C prior to reconstitution in PBS.

### *In vitro* immunological assays

#### SDS-PAGE gel electrophoresis and immunoblotting

In addition to the immunogens (the three recombinantly expressed SVSP toxins and corresponding snake venoms), an additional ten snake venoms (see Supplementary Table 1 for details), sourced from a wide geographical and taxonomic diversity of snake species known to cause potent haemotoxicity in snakebite victims, were also used for SDS-PAGE gel electrophoresis and western blotting experiments. These venoms were sourced from animals maintained under controlled environmental and dietary conditions in the UK Home Office licensed and inspected CSRI herpetarium at the Liverpool School of Tropical Medicine or from the CSRI historical venom collection. Lyophilised venoms were stored at 4 °C and reconstituted with phosphate-buffered saline (PBS, pH 7.4) to 1 mg/ml prior to use.

For gel electrophoresis, 15 well 15% SDS-PAGE gels were hand-cast, using the following approach: resolving gel, 3.75 ml H_2_O, 2.5 ml 1.5 M Tris pH 8.8, 3.75 ml 40 % bis-acrylamide, 100 µl 10 % SDS, 60 µl 10 % ammonium persulfate (APS) and 7 µl tetramethylethylenediamine (TEMED); stacking gel, 2.5 ml H_2_O, 1 ml 1 M Tris pH 6.8, 350 µl 40 % bis-acrylamide, 30 µl 10 % APS and 5 µl TEMED. Next, 10 µl of each recombinant toxin or venom (1 mg/ml) was mixed 1:1 volume/volume with reducing buffer (2 x PLOB; 3.55 µl H_2_O, 1.25 ml 0.5 M Tris pH 6.8, 2.50 ml glycerol, 2.0 ml 10 % SDS, 1.50 ml saturated bromophenol blue and 150 µl *β*-mercaptoethanol) and heated for 10-15 minutes at 100 °C. Thereafter, 10 µl was loaded onto the gel, alongside 5 µl of broad molecular weight protein marker (Broad Range Molecular Marker, Promega), and the samples were run at 200 volts for 55 minutes using a Mini-PROTEAN Electrophoresis System (Bio-Rad). The resulting gels were then stained at a final concentration of 0.1% (w/v) Coomassie blue R350 (0.4 g of Coomassie blue R350 in 200 mL of 40% [v/v] methanol in H_2_O, 10% (v/v) glacial acetic acid and 20% (v/v) methanol) overnight at room temperature. Gels were destained (4:1:5 methanol:glacial acetic acid:H2O) for at least 120 minutes at room temperature and images captured using a Bio-Rad Gel Doc EZ Gel Documentation System.

For immunoblotting experiments, SDS-PAGE gel electrophoresis was performed as described above, except with the use of prestained protein marker (10-225 kDa) (Thermo-Fisher Scientific, UK) and, instead of Coomassie staining, proteins in the gel were transferred onto 0.2 µm nitrocellulose membranes using a Trans-Blot Turbo Transfer System (Bio-Rad). Following confirmation of the transfer by reversible Ponceau S staining, membranes were blocked for non-specific binding using 5% non-fat dried milk in TBST (0.15 M NaCl; 0.01 M Tris-HCl, pH 8.5; 1% Tween 20), and left overnight at 4 °C on a rocker set at slow speed. Subsequently, blots were washed three times over 15 minutes with TBST, before the addition of primary antibodies (anti-ancrod, anti-batroxobin, anti-RVV-V, anti-toxin mix, anti-venom mix, and normal mouse control) diluted 1:5,000 in 5% non-fat dried milk in TBST for two hours at room temperature. The immunoblots were then washed in triplicate with TBST as described above and incubated for two hours at room temperature with 50 ml of horseradish peroxidase-conjugated Rabbit anti-mouse IgG (Sigma-Aldrich, UK) diluted 1:2,000 in PBS. The immunoblots were then rewashed with TBST and developed by adding DAB substrate (50 mg 3,3’- 222 diaminobenzidine, 100 ml PBS and 0.024% hydrogen peroxide, Sigma-Aldrich, UK) by placing the membrane into the substrate for 30 seconds before washing with deionised water.

### SDS-PAGE gel electrophoresis of deglycosylated recombinant toxins

To analyse the changes in molecular weight of glycosylated/deglycosylated recombinant toxins, we treated each recombinant protein with PNGase F. Briefly, recombinant toxins (14 μg) were mixed with 2 μl of 10X Denaturing Buffer (5% SDS, 400 mM Dithiothreitol [DTT]; Thermo-Fisher Scientific, UK) and ddH_2_0 to make a 20 μl total reaction volume, before incubation at 100 °C for five minutes. Next, 4 μl of 10X G7 Reaction Buffer (500 mM Sodium Phosphate pH 7.5), 4 µl of 10% triton™ X-100 (t- Octylphenoxypolyethoxyethanol; Sigma-Aldrich, UK), 1 μl PNGase F (New England Biolabs, UK) and 1 μl ddH_2_0 were added and samples left to digest overnight at 37 °C. Reaction products were then analysed by reduced SDS-PAGE gel electrophoresis as described above, with 40 µl 2 x PLOB reduction buffer and 60 µl PBS added to the 30 µl of each deglycosylated sample.

### End-point titration ELISA

Microtiter 96 well ELISA plates (Thermo-Fisher Scientific, UK) were coated with coating buffer (100 mM carbonate/Bicarbonate buffer, pH 9.6) containing 100 ng per well of each immunogen and incubated overnight at 4 °C. Plates were then washed three times with TBST before adding 5% non-fat milk in 100 ml TBST. Following incubation at room temperature for two hours, the plates were washed another three times with TBST. Next, 120 µl of the primary antibodies diluted 1:5,000 in 5% non-fat dried milk in TBST were added to the plate in duplicate at an initial dilution of 1:50 in 5% non-fat milk in TBST, followed by five-fold serial dilutions across the plate, and incubation at 4 °C overnight. The plates were then rewashed with TBST and incubated for two hours at room temperature with appropriate secondary antibodies (horseradish peroxidase-conjugated rabbit anti-mouse IgG, Sigma-Aldrich, UK), diluted to 1:2,000 in PBS. The plates were then rewashed with TBST before the addition of substrate (0.2% 2,2/-azino-bis (2-ethylbenzthiazoline-6-sulphonic acid) in citrate buffer (0.5 M, pH 4.0) containing 0.015% hydrogen peroxide, Sigma-Aldrich, UK). Plates were gently mixed and incubated at room temperature for 15 minutes before the signal was read spectrophotometrically at 405 nm on a FLUOstar Omega microplate reader.

### *In vitro* functional assays

#### Fibrinogenolytic activity

To assess the fibrinogenolytic activity of the immunogens and their inhibition by the resulting experimental antivenoms, we used a degradation SDS-PAGE gel electrophoresis approach using human plasma fibrinogen (Sigma-Aldrich, UK). The recombinant toxin immunogens (7 μg, 1 mg/ml) and various crude venoms (7 μg, 0.3 mg/ml) were incubated with human fibrinogen (3 µg, 2.5 mg/ml, Sigma-Aldrich, UK) for two hours at 37 °C. All immunogen samples were also used in neutralisation experiments following a pre-incubation step at 37 °C for 15 minutes with 0.5 µg (1 mg/ml) of the corresponding toxin-specific experimental antivenom (i.e., for ancrod and *C. rhodostoma* venom, anti-ancrod was used). Next, ten well 8% SDS-PAGE gels were hand-cast using the following approach: 10 ml resolving gel (4.7 ml H_2_O, 2.5 ml Tris pH 8.8 (1.5 M), 2.7 ml 30% bis-acrylamide, 50 µl 20% SDS, 100 µl 10% APS and 7 µl TEMED); 4 ml of 4% stacking gel (2.7 ml H_2_O, 0.5 ml Tris pH 6.8 (1 M), 800 µl 30% bis-acrylamide, 20 µl 20% SDS, 40 µl 10% APS and 4 µl TEMED). Thereafter, 10 µl of each sample was loaded on the gel and analysed under reducing conditions as described earlier. The negative control consisted of human fibrinogen only (3 µg, 2.5 mg/ml, Sigma-Aldrich, UK).

### Blood sample collection and reagents for clotting profiling

Blood samples for clotting profiling experiments were obtained according to ethically approved protocols (LSTM research tissue bank, REC ref. 11/H1002/9) from consenting healthy volunteers who confirmed they had not taken any anticoagulant treatments for at least three months prior to blood collection. Blood samples were collected in tubes containing acid citrate dextrose adenine (ACD-A) as an anticoagulant. To prepare Fresh Frozen Plasma (FFP) fresh blood samples were centrifuged at 2500 × g at 20-25 °C for 10 minutes, and the supernatant was retained and stored at -80 °C until use. We used commercially sourced samples (Diagnostic Reagents Ltd, UK) to serve as quality controls for the various reagents employed in the methods described below. To this end, normal and abnormal test controls were implemented for the experiments described below prior to experimentation. These consisted of 1) Diagen control plasma for fibrinogen: normal (RCPN070) and abnormal (RCPA080) and 2) control plasma for aPTT and PT: normal (IQCN130), abnormal 1 (Mild, IQCM140) and abnormal 2 (Severe, IQCS150). All QC samples were aliquoted into 500 µl, stored at -80°C, and defrosted in a water bath for five minutes at 37 °C immediately prior to use. To act as positive controls for the experimental antivenoms generated here, we used commercially available antivenoms, specifically the Thai Red Cross monovalent equine antivenoms directed against the Malayan Pit Viper *C. rhodostoma* (CRAV; Lot #CR00316, expiry date 06/2021) and the Russell’s viper *D. siamensis* (DSAV; Lot #WR00117, expiry date 11/2022) as controls for *C. rhodostoma* and *D. russelli* venom respectively, and the Instituto Butantan polyvalent equine SORO antibotropico/crotalico antivenom (Lot #1012308, expiry date: 2013) for *B. atrox*. Antivenoms were reconstituted with pharmaceutical grade water supplied by the manufacturer, protein concentrations measured using a Nanodrop (Thermo-Fisher Scientific, UK), and then stored short term at 4 °C until use. Normal horse IgG (1 mg/mL; Bio-Rad, UK) and normal mouse IgG (1 mg/ml, as above) were used as negative controls for the commercial antivenoms and experimental antivenoms, respectively, throughout.

### Activated Partial Thromboplastin Time (aPTT)

To measure the inhibitory capability of the experimental antivenoms against recombinant toxins acting on the intrinsic and common coagulation pathways, we quantified differences in the activated Partial Thromboplastin Time (aPTT) between toxin and venom samples in their presence and absence. To do so, 50 µL of Micronised Silica/Platelet Substitute Mixture (Diagnostic Reagents Ltd, UK) was placed in glass test tubes (10 × 75 mm) in a water bath at 37 °C and incubated for 60-120 seconds. Next, 50 µl of either FFP was spiked with 0.6 ng of ancrod, batroxobin, RVV-V, *C. rhodostoma* venom, *B. atrox* venom or *D. russelii* venom, or saline solution as the negative control. All immunogen samples were also co-incubated at 37 °C for 15 minutes with 0.5 µg of either: (i) the corresponding toxin-specific experimental antivenom (i.e., for ancrod, anti-ancrod was used), (ii) the anti-toxin mix experimental antivenom, (iii) the anti-venom mix experimental antivenom, (iv) the corresponding commercial antivenom (i.e., for ancrod the Malayan pit viper antivenom was used) and (iv) normal mouse IgG as a non-immunised control. Thereafter, samples were added to the glass test tubes and gently tilted at regular intervals for precisely five minutes at 37 °C. Finally, 50 µL of 25 mM calcium chloride (pre-incubated at 37°C) was added to each tube, and the tube was gently tilted until the resulting clot time was recorded. All experiments were performed in duplicate.

### Prothrombin Time

To measure the inhibitory capability of the experimental antivenoms against recombinant toxins acting on the extrinsic coagulation pathway, we quantified differences in the prothrombin time (PT) between toxin and venom samples in the presence and absence of the different antivenoms. Measurements of PT were undertaken by first adding 100 µl of Calcium Rabbit Brain Thromboplastin (Diagnostic Reagents Ltd, UK) to a glass test tube (10 × 75 mm) and incubating at 37 °C for 60-120 seconds in a water bath. Next, 50 µl of FFP was spiked with 0.6 ng of ancrod, batroxobin, RVV-V, *C. rhodostoma* venom, *B. atrox* venom or *D. russelii* venom, or saline solution as the negative control. All immunogen samples were also co-incubated at 37 °C for 15 minutes with 0.5 µg of either: (i) the corresponding toxin-specific experimental antivenom (i.e., for ancrod, anti-ancrod was used), (ii) the anti-toxin mix experimental antivenom, (iii) the anti-venom mix experimental antivenom, (iv) the corresponding commercial antivenom (i.e., for ancrod the Malayan pit viper antivenom was used) and (iv) normal mouse IgG as a non-immunised control. Next, samples were introduced to the thromboplastin and time measurements were commenced, with tubes gently tilted at regular intervals (returning to the water bath between tilting), and the time for the formation of a clot recorded. All experiments were performed in duplicate.

### Fibrinogen consumption via the Clauss method

The Clauss method is a quantitative, clot-based assay that measures the ability of thrombin to convert fibrinogen to a fibrin clot, followed by manual time measurements of clotting (45). Here we applied this method in recombinant toxin- and venom-spiking experiments to assess the inhibitory capability of the experimental antivenoms against the depletion of fibrinogen. Twenty microlitres of FFP was spiked with 0.6 ng of either: ancrod, batroxobin, RVV-V, *C. rhodostoma* venom, *B. atrox* venom or *D. russelii* venom, or 0.9 % saline solution as the negative control. All immunogen samples were also co-incubated at 37 °C for 15 minutes with 0.5 µg of either: (i) the corresponding toxin-specific experimental antivenom (i.e., for ancrod the anti-ancrod was used), (ii) the anti-toxin mix experimental antivenom, (iii) the anti-venom mix experimental antivenom, (iv) the corresponding commercial antivenom (i.e., for ancrod the Malayan pit viper antivenom was used) or (iv) normal mouse IgG as a non-immunised control. Samples were then diluted tenfold with 0.02 M imidazole buffer (pH 7.35), transferred to glass test tubes (10 × 75 mm) and warmed at 37 °C for 120 seconds. Thereafter, 100 µl of thrombin reagent (20 units/ml; Diagnostic Reagents Ltd, UK) was added, and time measurements commenced. Tubes were gently tilted at regular intervals (returning to the water bath between tilting), and the time for the formation of a clot was recorded. To calculate the fibrinogen concentration (g/L) from the clotting time, calibration curve analysis was used. All experiments were performed in duplicate.

## Results

### Expression, purification, and evaluation of recombinantly expressed SVSP toxins

The isolated DNA of the three selected recombinant SVSPs toxins were transformed using TOP10 *E. coli* competent cells, transfected into mammalian HEK293F cells, and the resulting expressed protein purified using hydrophobic affinity chromatography. Reduced 10% SDS-PAGE gel electrophoretic profiles of the resulting eluted recombinant toxins revealed protein bands corresponding with the expected molecular mass range of SVSP toxins, and demonstrated a single sharp protein band for both RVV-V and batroxobin and a single band extending over several kilodaltons (≈ 35-45 kDa) for ancrod, suggesting the production of several different glycoforms (see right hand columns of Figure 1A-C). Treatment of each toxin with the deglycosylating agent PNGase F resulted in findings supporting this assertion, with molecular weight shifts observed for each protein, including the resolution of the molecular weight range for ancrod resolving into a single protein band (Supplementary Figure 2). To test whether the three recombinant toxins were correctly folded, we next used degradation SDS-PAGE gel electrophoresis to assess their fibrinogenolytic activity. Our findings revealed that recombinantly expressed ancrod, batroxobin and RVV-V were all functionally active, as evidenced by effective cleavage of the *α* chain of fibrinogen by all three proteins, alongside additional partial cleavage of the *β* chain by batroxobin (Figure 1A-C).

**Figure 1.**
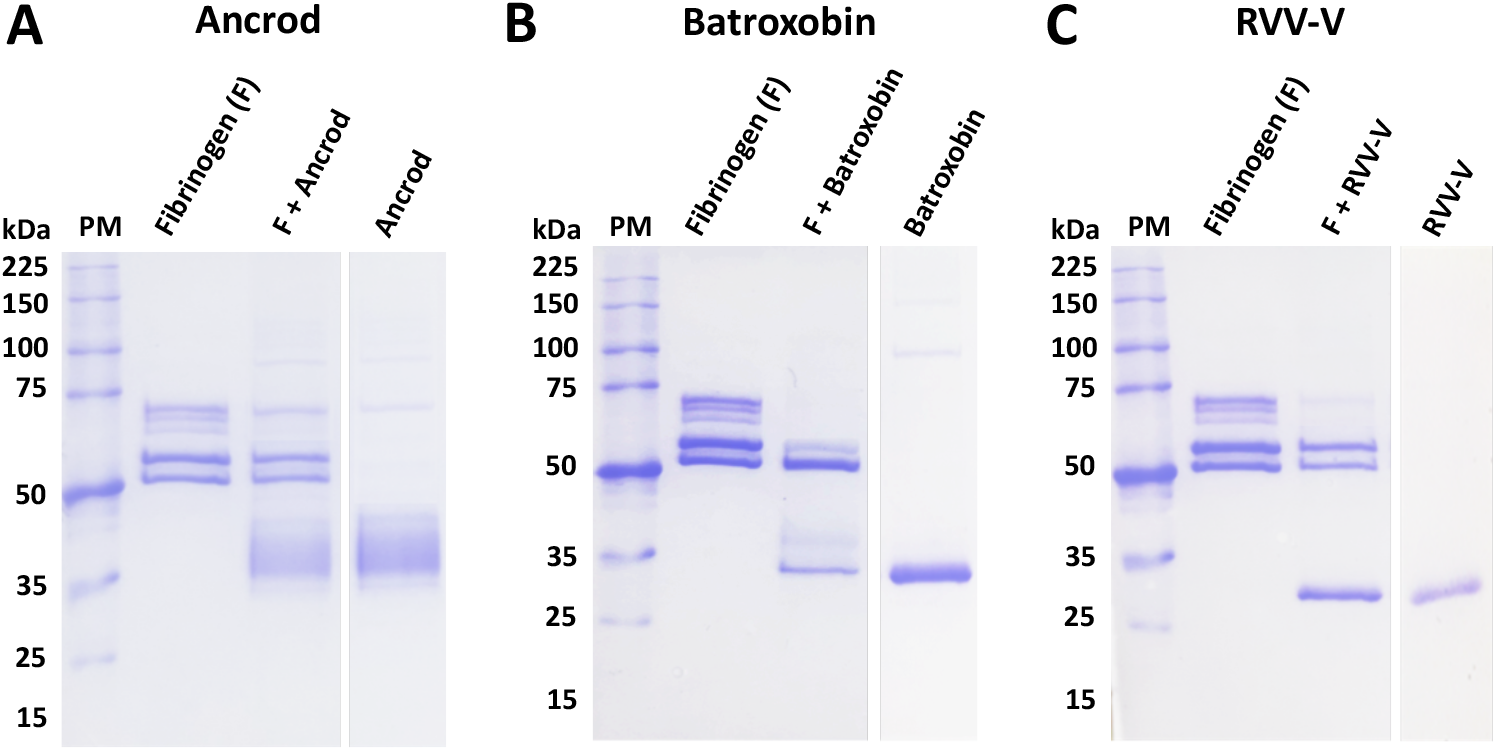
The protein profiles of the purified recombinant toxins and their activity on human fibrinogen. Degradation SDS-PAGE gel electrophoretic profiles (reduced conditions, 8% gel) displaying the fibrinogenolytic activity of the recombinant toxins (7 μg) following their incubation with human fibrinogen (3 μg) at 37 °C for 120 min: **(A)** ancrod, **(B)** batroxobin and **(C)** RVV-V. PM represents the molecular weight protein marker.

### Monitoring seroconversion to the toxin immunogens

The protein concentrations recovered for each recombinantly expressed toxin following purification resulted in sufficient yields (∿0.7 mg/ml) for murine immunisation experiments. Over 14 weeks, different groups of mice received multiple immunising doses of each recombinant toxin, alongside a group receiving a mixture of all three toxins (in a 1:1:1 ratio), and a control group that received a native venom mixture. To quantify antibody binding levels to the immunogens, serum samples were collected at weeks 3, 6, 10 and 14 (terminal sample) and assessed via ELISA. The mice responded to the recombinant immunogens in a variable manner, though gradual increases in antibody binding levels were typically observed following primary immunisation until the end of the experiment (Figure 2). The exception to this was the responses observed in the group immunised with batroxobin, which showed lower binding levels until week eight (Figure 2). The resulting titres of specific antibodies were highest in the groups immunised with ancrod (Figure 2A) and the 1:1:1 mixture of the three recombinant toxins (Figure 2D). In addition to monitoring responses against the immunogens, we also quantified time-course immunological responses to the three crude venoms from which the recombinant toxins are derived. As anticipated, binding levels were reduced compared with the immunogens, likely due to SVSPs toxins only making up a proportion of the crude venoms compared with the recombinant toxin immunogens, though binding levels of the anti-batroxobin sera against *B. atrox* venom remained very low even following the third and fourth immunisation (Figure 2F). However, binding levels of the anti-ancrod and anti-RVV-V sera against the corresponding venoms (i.e. *C. rhodostoma* and *D. russelii* venom, respectively) provided clear evidence that these two recombinant toxins stimulated the production of antibodies by the end of the experimental time course that recognised native venom proteins (Figures 2E and 2G). High levels of immunological binding were observed between the positive control experimental sera (i.e. the “anti-venom mix”) and the three crude snake venoms mixed and used as the immunogen, with maximal binding effectively achieved after only four weeks (Figure 2H), unlike that observed with the recombinant proteins. Nonetheless, as anticipated, mice serum samples collected at the end of the immunisation schedule (14 weeks) exhibited the highest antibody binding titres across all groups. These samples were subjected to IgG purification to produce the experimental antivenoms used for all downstream analyses.

**Figure 2.**
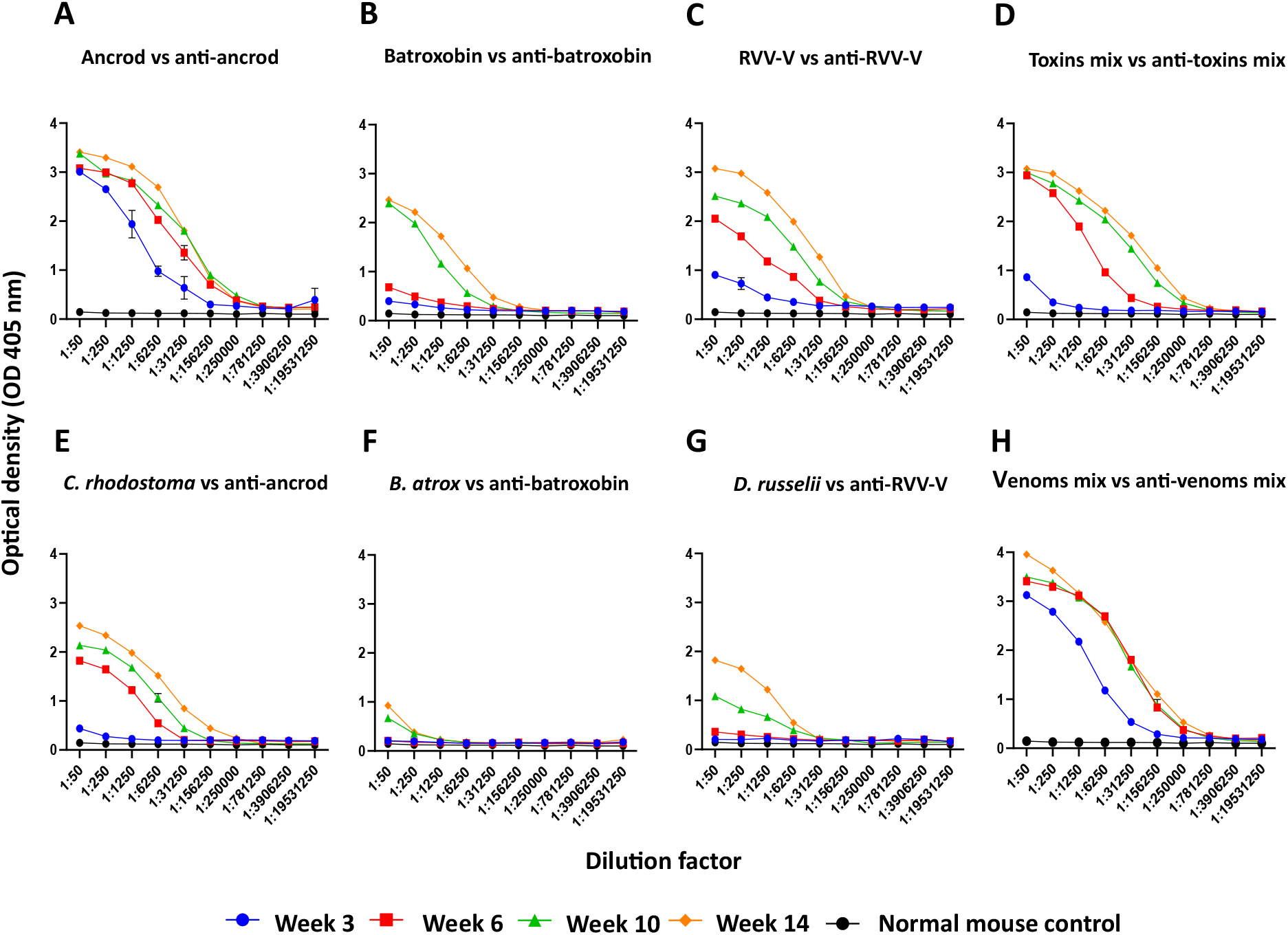
Time-course analysis of the immunological cross-reactivity of pooled sera to the recombinant toxins and crude venoms used as immunogens over 14 weeks of murine immunisation. Responses of anti-ancrod pooled mice sera against ancrod; **(B)** anti-batroxobin sera against batroxobin; **(C)** anti-RVV-V sera against RVV-V; **(D)** anti-toxin mix sera against a 1:1:1 mixture of the three recombinant toxins; **(E)** anti-ancrod sera against *C. rhodostoma* venom; **(F)** anti-batroxobin sera against *B. atrox* venom; **(G)** anti-RVV-V sera against *D. russelii* venom; **(H)** anti-venom mix sera (positive control) against a 1:1:1 mixture of the three crude venoms. Data is shown for sera collected at weeks 3, 6, 10 and 14 (end of the experiment) of the immunisation time course. Non-immunised mouse sera (“normal mice control”) was used as a negative control. All mice serum samples were standardised to 1:50, then diluted fivefold. Data points represent means of duplicate readings, and error bars represent the standard deviation (SD).

### Quantifying and visualising immunological cross-reactivity

We used endpoint ELISA (EPT-ELISA) experiments to quantify the binding levels detected between the resulting experimental IgG antivenoms and the toxins and venoms used as immunogens. Comparisons of the binding profiles revealed that responses to ancrod, RVV-V and the mixture of the three recombinant toxins were highly consistent across the different experiential antivenoms generated and stimulated the highest binding titres (Figure 3A-D). Notably, binding levels to each recombinant toxin by the corresponding experimental antivenoms were higher than the positive control antivenom (i.e. generated via immunisation with the three crude venoms), presumably as the result of the majority of antibodies being directed towards specific SVSP immunogens, rather than the broad diversity of toxins present in the crude venoms (Figure 3). Despite these promising findings, the anti-batroxobin antivenom generally underperformed (consistent with serology data shown in Figure 2) and exhibited binding titres considerably lower than the other experimental antivenoms against each immunogen, with the exception of cross-reactivity to recombinant batroxobin (Figure 3A-D). The immunological responses to crude venoms by the experimental antivenoms were also variable. Unsurprisingly, the highest binding levels were observed with the positive control antivenom (i.e. generated using the three venom mix as an immunogen). While the anti-ancrod, anti-RVV-V and anti-toxin mix antivenoms all exhibited moderate binding against *C. rhodostoma* venom and the three-venom mixture (Figure 3E and 3H), responses to *B. atrox* and *D. russelii* venom were very low (Figure 3F and 3G), perhaps suggesting low SVSP abundance in these venoms.

**Figure 3.**
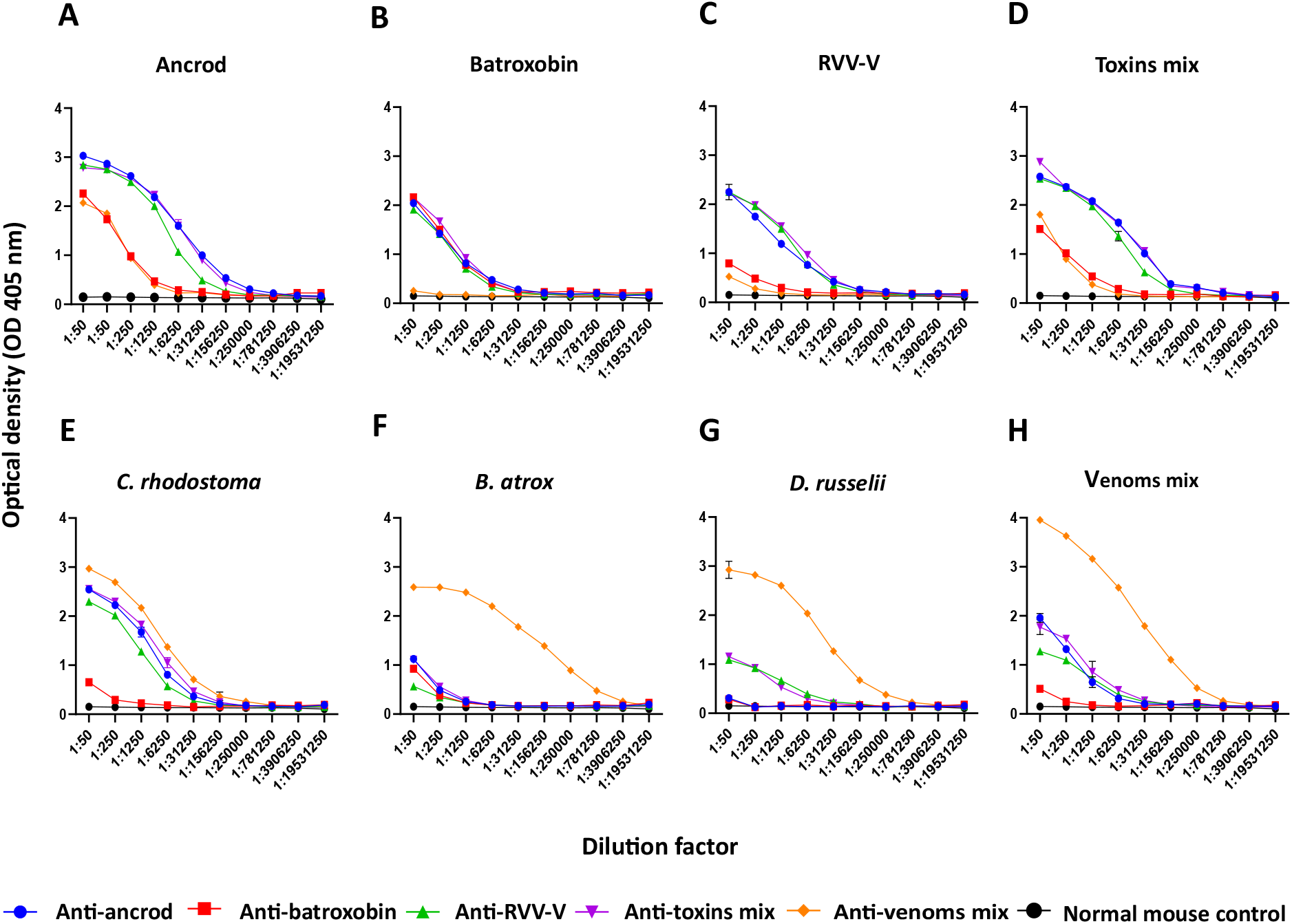
EPT-ELISA analyses of immunological binding between the experimental antivenoms and the recombinant toxins and crude venoms used as immunogens. Each resulting experimental antivenom (anti-ancrod, anti-batroxobin, anti-RVV-V, anti-toxin mix and anti-venom mix) is coloured differently, and their binding to the various toxin and venom immunogens are displayed in different panels, alongside data obtained with the normal mouse control negative control. Data shown represents binding levels to: **(A)** ancrod, **(B)** batroxobin, **(C)** RVV-V, **(D)** a mixture of the three recombinant toxins, **(E)** *C. rhodostoma* venom, **(F)** *B. atrox* venom, **(G)** *D. russelii* venom, and **(H)** a mixture of the three venoms. Each antivenom sample was serially diluted fivefold in duplicate, with data points representing means of duplicate readings, and error bars representing standard deviations (SD).

Next, reduced SDS-PAGE gel electrophoresis and western blotting experiments were performed to visualise the immunological recognition of the experimental antivenoms to each of the toxins and venoms used as immunogens (Figure 4). Each of the experimental antivenoms generated with recombinant toxin immunogens exhibited immunological recognition of each of the three recombinant toxins, irrespective of which was used to generate the antibodies (Figure 4B-D). Further, both the anti-ancrod and anti-RVV-V antivenoms displayed recognition of various toxins found in the crude venoms of *C. rhodostoma, B. atrox* and *D. russelii* in a highly comparable manner (Figure 4A and 4C), though recognition by the anti-batroxobin antivenom was considerably reduced (Figure 4B), consistent with the earlier ELISA experiments (Figure 3). When tested against a broader panel of crude snake venoms, antibodies generated against the recombinant toxins showed immunological cross-reactivity with proteins exhibiting molecular weights consistent with SVSP toxins present in other pitvipers (*Bothrops* and *Crotalus* spp.) (Supplementary Figure 3), and these findings were also consistent with EPT-ELISA binding levels being increased against these venoms, except in the case of the anti-batroxobin experimental antivenom (Supplementary Figure 4). The positive control antivenom generated against the mixture of the three crude venoms exhibited a distinct binding pattern, characterised by broad binding against the diversity of toxins found in each of the crude venoms (both those used as immunogens and others), but with only noticeable recognition of ancrod out of the three toxins recombinantly expressed in this study (Figure 4F and Supplementary Figures 3 and 4).

**Figure 4.**
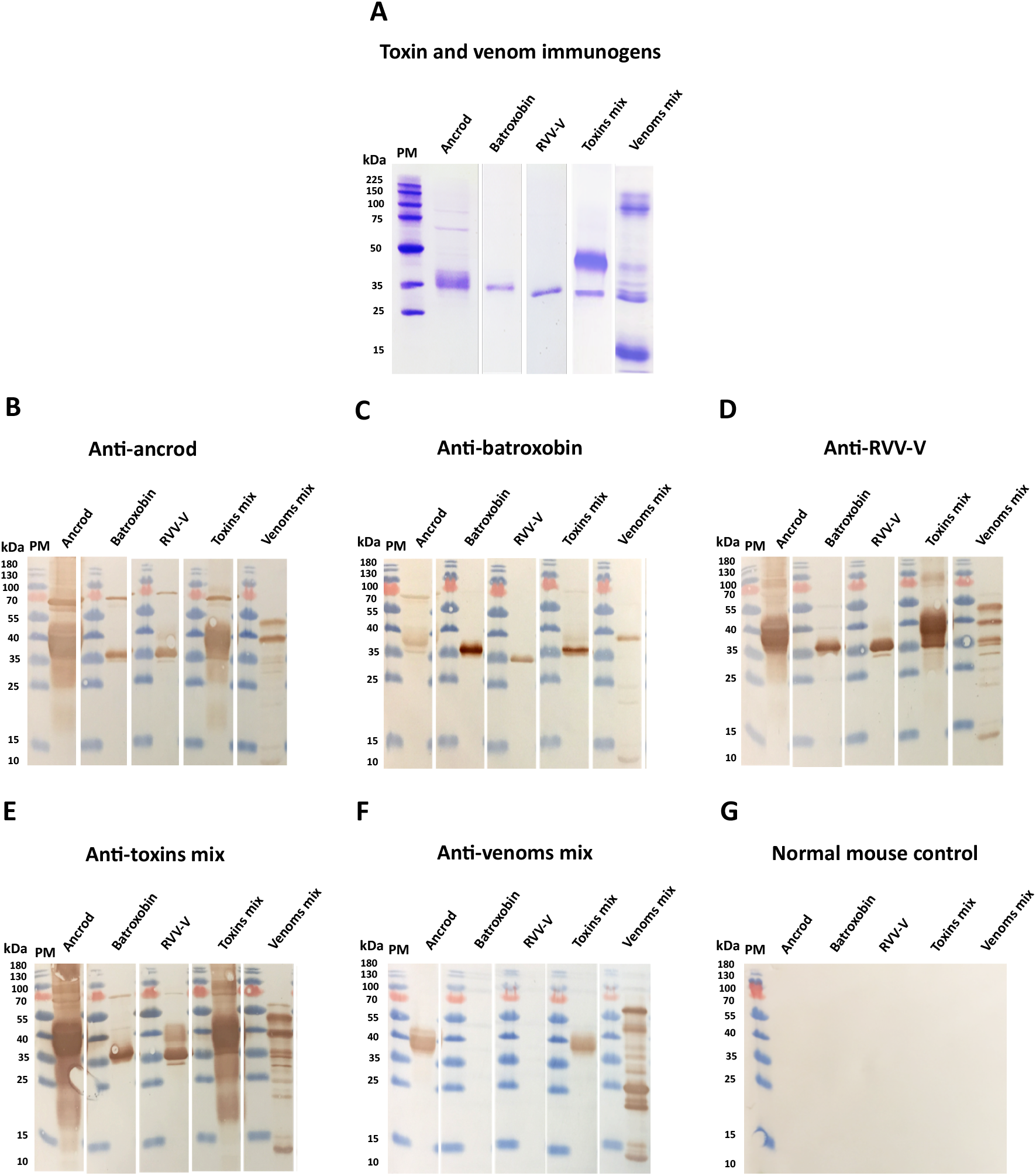
Immunological recognition of the toxin and venom immunogens by the different experimental antivenoms. **(A)** Reduced 15% SDS-PAGE gel electrophoresis and Coomassie blue staining was used to visualise the toxin and venom immunogens (ancrod, batroxobin, RVV-V, a 1:1:1 mix of these three toxins, and a 1:1:1 mix of *C. rhodostoma, B. atrox* and *D. russelii* venoms). The same venom samples were transferred to nitrocellulose membranes for immunoblotting experiments and incubated with 1:5,000 dilutions of primary antibodies of each of the experimental antivenoms, specifically: **(B)** anti-ancrod, **(C)** anti-batroxobin, **(D)** anti-RVV-V, **(E)** anti-toxin mix, **(F)** anti-venom mix (as positive control) and **(G)** normal mouse IgG (as negative control). PM indicates protein marker, and note that different molecular weight markers were used for SDS-PAGE and immunoblotting experiments.

### Inhibition of fibrinogenolytic activity

Many SVSP toxins exhibit fibrinogenolytic activity similar to human thrombin (i.e., TLEs). While thrombin can convert fibrinogen to fibrin via cleavage of the *α* and *β* chains (46), venom TLEs exert variable specificities and can cleave either both the *α* and *β* chains, or preferentially cleave either the *α* or *β* chains, of fibrinogen (47). During validation of protein expression, we demonstrated that recombinant ancrod, batroxobin and RVV-V all cleaved the *α* chain of fibrinogen (Figure 1). Co-incubation of these recombinant toxins with their corresponding experimental antivenoms resulted in inhibition of fibrinogenoylsis, as evidenced by visualisation (i.e. restoration vs toxin only control) of the *α* chain (Figures 5A-C), and demonstrating that the immunological cross-reactivity described above can also confer inhibition of toxin function. Unsurprisingly, crude venoms from *C. rhodostoma, B. atrox* and *D. russelii* were also able to cleave the *α* chain of fibrinogen, though *D. russelii* was the least potent in this regard, and *C. rhodostoma* venom additionally cleaved the *β* chain (Supplementary Figure 5). Although inhibition experiments with the corresponding experimental antivenoms did not result in complete inhibition of venom activity, likely due to distinct non-SVSP toxins also contributing, in each case the experimental antivenoms did reduce the extent of venom-induced fibrinogenolysis observed (Supplementary Figure 5). Comparisons with the normal mouse control, where no inhibition was observed (Supplementary Figure 6), demonstrated that this effect was the result of antibody specificities stimulated by the recombinant toxin immunogens.

**Figure 5.**
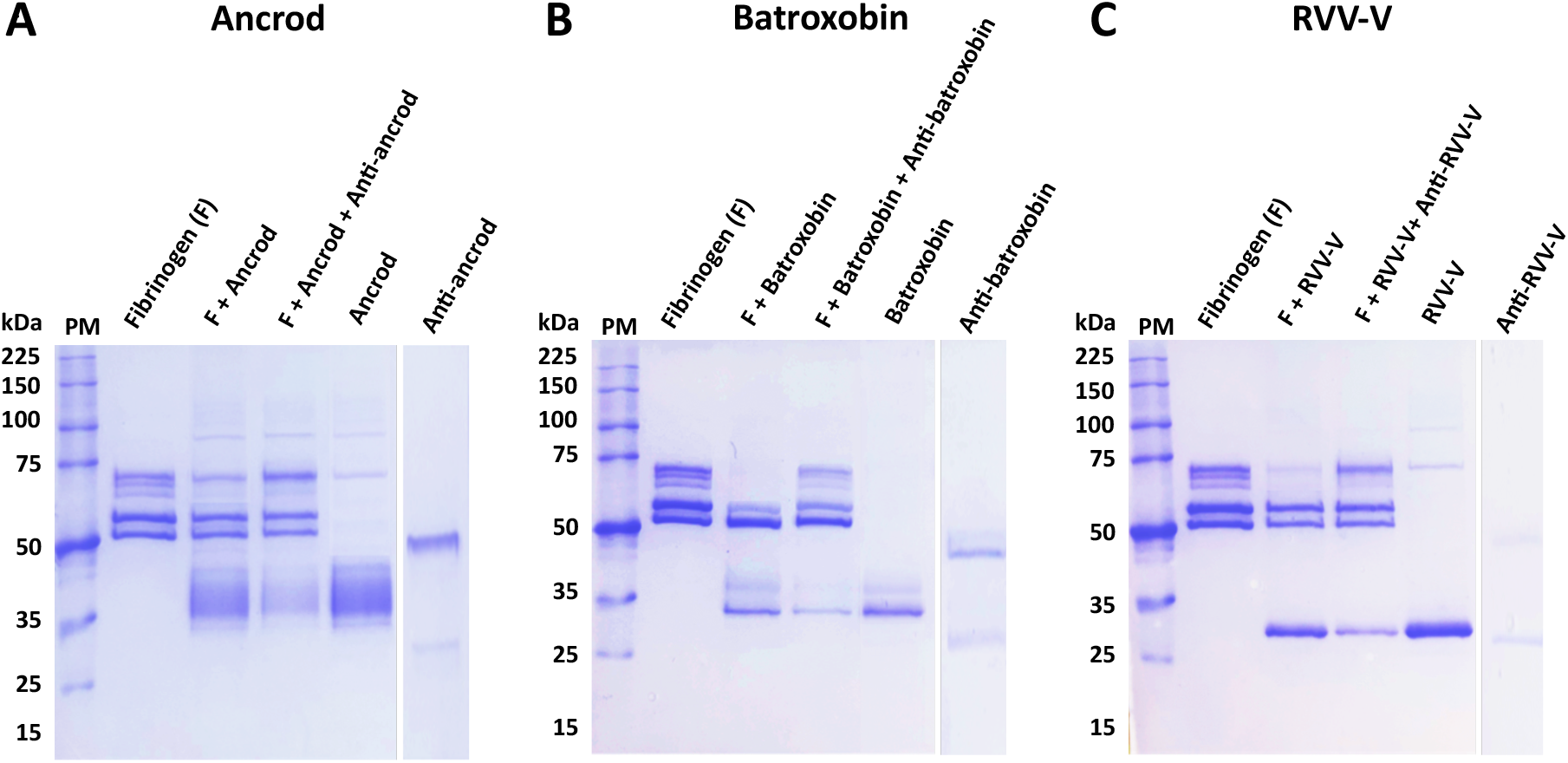
Experimental antivenoms directed against recombinant SVSP toxins inhibit their fibrinogenolytic activity. Degradation SDS-PAGE gel electrophoretic profiles are displayed following the incubation of various samples at 37 °C for 120 min. Panels show different data obtained with the different recombinant toxins: **(A)** Ancrod, **(B)** Batroxobin and **(C)** RVV-V. For each, the following layout was used: Lane 1, protein marker (PM); Lane 2, human fibrinogen (3 μg); Lane 3, fibrinogen + recombinant toxin (7 μg, ancrod, batroxobin or RVV-V); Lane 4, fibrinogen + recombinant toxin + experimental antivenom (0.5 μg, anti-ancrod, anti-batroxobin or anti-RVV-V); Lane 5, recombinant toxin only; Lane 6, specific experimental antivenom only.

### Inhibition of coagulation disturbances

To assess the inhibitory capability of the experimental antivenoms against recombinant toxins and crude venoms acting more broadly on components of the blood clotting cascade, we next quantified various coagulation parameters. First, using FFP, we measured the PT and aPTT stimulated by the recombinant toxins and corresponding crude venoms in the presence and absence of our experimental antivenoms, and using commercial antivenoms as controls. The PT measures clotting dictated by the extrinsic and common coagulation cascades, and a prolonged PT can result from an absence or deficiency of one or more of Factors X, VII, V, II or I (fibrinogen). Our results showed that each of the recombinant toxins and corresponding crude venoms substantially prolonged the PT, with the crude venoms resulting in increased clotting time delays over each of the corresponding recombinant toxins (Figure 6), likely due to additional toxins contributing to the overall effect on the coagulation cascade. In terms of inhibition, although the normal mouse control (i.e. containing non-specific antibodies) noticeably reduced the venom/toxin-induced PT prolongation observed in these experiments, suggesting considerable non-specific inhibitory effects, the experimental antivenoms directed against specific recombinantly expressed SVSP toxins exhibited superior inhibitory profiles. Indeed, at the doses of antivenom tested, the anti-toxin experimental antivenoms consistently produced the greatest levels of venom inhibition, exceeding those of the experimental anti-toxin mix anti-venom mix antivenoms and also the specific commercial antivenoms, and generally reduced toxin-and venom-induced PT prolongations reduced to near control levels (Figure 6). Contrastingly, none of the recombinant toxins or crude venoms under study had an effect on the aPTT, which measures clotting dictated by the intrinsic and common coagulation cascades, and thus inhibitory effects could not be measured (Supplementary Figure 7).

**Figure 6.**
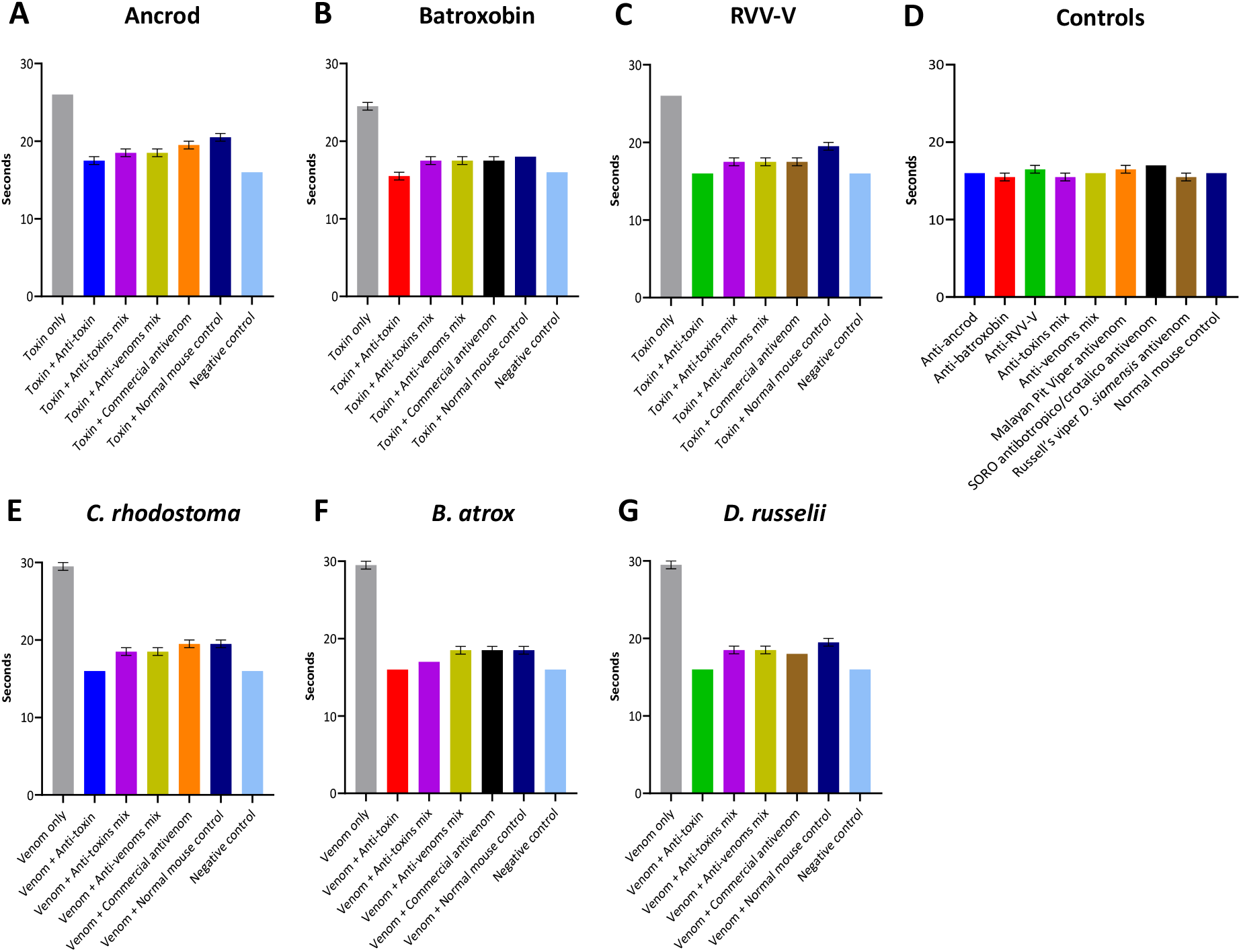
Inhibition of coagulation disturbances defined by the prothrombin time (PT). The assay measured the combined effect of the clotting factors of the extrinsic and common coagulation pathways (in seconds) in the presence of the recombinant toxins/crude venoms, and their recovery effect by adding specific experimental antivenoms/specific commercial antivenoms, incubated with FFP. **A)** Ancrod, **B)** Batroxobin, **C)** RVV-V, **D)** the various controls, **E)** *C. rhodostoma* venom, **F)** *B. atrox* venom and **G)** *D. russelii* venom. For each toxin/venom, “homologous” antivenom combinations were used (e.g. for ancrod the anti-ancrod antivenom was used as the anti-toxin antivenom, and the Malayan pit viper antivenom was used as the commercial antivenom). Each experimental antivenom alone, each commercial antivenom alone and normal mouse control alone were used as negative controls. Error bars represent the standard deviation (SD) of duplicate measurements.

The clotting time of diluted plasma with a standard concentration of thrombin is inversely related to the fibrinogen concentration, and at low fibrinogen concentrations, the reaction rate is, therefore, a function of fibrinogen concentration (45). To further assess the inhibitory capabilities of the experimental antivenoms against the fibrinogenolytic activity of the recombinant SVSP toxins and venoms used as immunogens, we quantified the consumption of fibrinogen using clotting time experiments with FFP. Quantification of resulting fibrinogen concentrations demonstrated that the three SVSP toxins (ancrod, batroxobin and RVV-V) and the three native snake venoms tested (*C. rhodostoma, B. atrox* and *D. russelii*) all dramatically reduced fibrinogen levels compared to the control (0.73-1.96 g/L vs 3.68 g/L, respectively) (Figure 7). Noticeably, none of the antivenoms tested, including the commercial antivenom controls, were able to restore fibrinogen levels to those of the control, irrespective of which recombinant toxin or venom was used. Broadly speaking, the anti-toxin mix and anti-venom mix experimental antivenoms exhibited highly comparable inhibitory responses, though both provided either highly comparable or modest additional reductions in fibrinogen depletion over the commercial antivenom control (Figure 7). However, these resulting fibrinogen levels were generally similar to, or slightly higher than, those obtained using the non-specific normal mouse control samples. Contrastingly, the toxin-specific experimental antivenoms (i.e. anti-ancrod, anti-batroxobin and anti-RVV-V) displayed clear evidence of protection against fibrinogen depletion, with the resulting fibrinogen concentrations recovered in all homologous toxin and venom combinations substantially higher than the corresponding toxin or venom only control values (2.23-2.56 g/L vs 0.73-1.96 g/L) (Figure 7).

**Figure 7.**
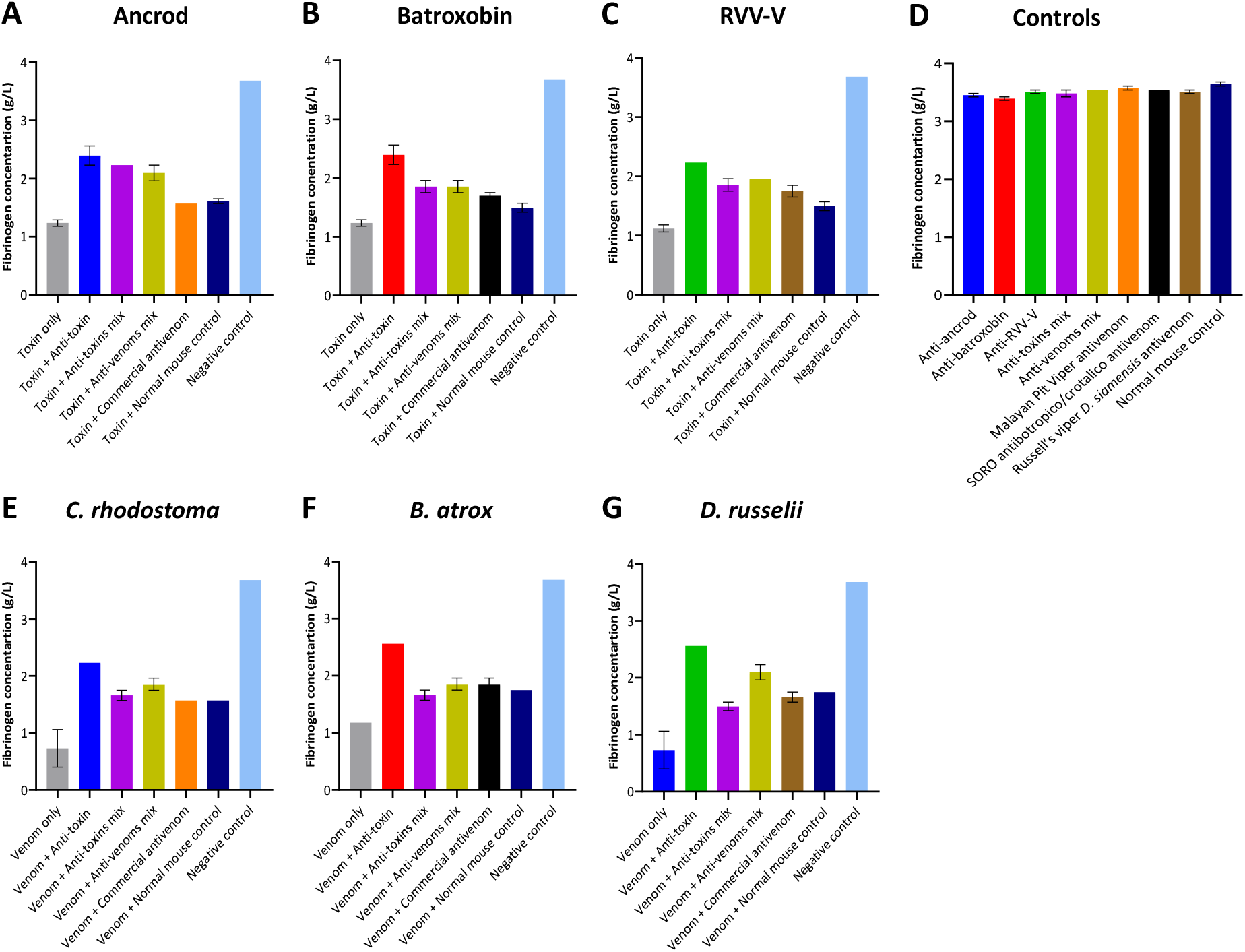
Quantification of fibrinogen concentrations following coincubation of recombinant toxins and crude venoms with the specific experimental and commercial antivenoms and human plasma. The assay uses an excess of thrombin to convert fibrinogen to fibrin in diluted citrated human fresh frozen plasma (FFP). The resulting fibrinogen concentration is shown for each recombinant toxin/crude venom against the specific experimental antivenoms incubated in FFP. Data shown represents the following immunogens: **A)** Ancrod, **B)** Batroxobin, **C)** RVV-V, **D)** the various controls, **E)** *C. rhodostoma* venom, **F)** *B. atrox* venom and **G)** *D. russelii* venom. For each toxin/venom, “homologous” antivenom combinations were used (e.g. for ancrod the anti-ancrod antivenom was used as the anti-toxin antivenom, and the Malayan pit viper antivenom was used as the commercial antivenom). Each experimental antivenom alone, each commercial antivenom alone and the normal mouse control alone were used as negative controls. Error bars represent the standard deviation (SD) of duplicate measurements.

## Discussion

Antivenom is the only specific treatment for snakebite envenoming, and though these therapeutics save countless lives each year, they have several limitations that need to be overcome to ensure effective, safe and affordable treatment is accessible for tropical snakebite victims. One of the main limitations with current antivenoms is that only about 10-20% of the antibodies present in them are specific to the toxins found in the venom immunogens (23, 27). Furthermore, of that 10-20%, a substantial proportion will be directed towards toxins that may not necessarily be of great importance for neutralisation, for example if they are of low toxicity or relevance for causing pathology in snakebite victims (48). Consequently, there have been many experimental attempts to improve conventional antivenoms by focusing the immune response towards the generation of polyclonal antibodies against key pathogenic toxins, rather than all toxins, via the use of immunogens distinct from crude venom. Examples of such strategies include the use of strings of linear epitopes (49), virus-like particles decorated in conserved epitopes (50), recombinantly expressed venom toxins (51) and recombinantly expressed consensus toxins (52).

In this study, we explored the potential utility of using recombinantly expressed toxins to generate polyclonal antibody responses directed against the haemotoxic SVSP toxin family. The SVSPs are enzymatic proteases that are common toxins in snake venom systems and have been studied for many decades due to their ability to interfere with haemostasis in a variety of ways (53). Many SVSPs act as thrombin-like enzymes (i.e. the TLEs) and during envenoming their fibrinogenolytic activity contributes towards VICC and the severity of haemotoxicity (8, 11), thus making them priority targets for neutralisation. In this study we applied a mammalian cell expression approach for SVSP immunogen production, rather using other expression systems or native toxin purification. We chose mammalian cells for expression because SVSPs are often glycosylated and have a number of disulfide bonds, which present obstacles to effectively refold the recombinant proteins as active forms from *E. coli-*expressed insoluble inclusion bodies, even under conditions intended to favour disulfide bond formation (35, 36). Such refolding steps are crucial but are frequently unsuccessful and/or time-consuming to optimise sufficient yield recovery, as reported in several previous snake venom toxin expression studies (31, 32, 54, 55). Consequently, here we used mammalian HEK293F cells for the expression of three pathogenically relevant SVSP toxins sourced from geographically diverse medically important viperid snakes, specifically ancrod from *C. rhodostoma*, batroxobin from *B. atrox* and RVV-V from *D. russelli*. Expression of each protein resulted in acceptable yields (≈ 0.7 mg from 1 ml culture), purity and, since many SVSPs are variably glycosylated (56, 57), evidence of glycosylation. As a measure of success of correct disulfide bond formation and native protein folding, we tested the functional activities of the toxins in haemotoxicity bioassays, which revealed each exhibited fibrinogenolytic activities, and dramatically decreased fibrinogen concentrations and prolonged prothrombin times when incubated with FFP.

To further explore the utility of recombinant toxins for the future development of snakebite antivenoms with more focused anti-toxin inhibitory profiles, we used the recombinantly expressed SVSPs as immunogens. The toxins elicited variable immune responses, though the resulting experimental murine antivenoms demonstrated promising cross-reactivity to both the recombinant SVSP immunogens and native toxins found in a variety of crude snake venoms. Although comparisons of the resulting binding profiles indicated that responses to ancrod, RVV-V, and the recombinant toxin mixture were strong and highly consistent across the different experimental antivenoms generated, the responses generated by recombinant batroxobin were considerably reduced, except against batroxobin itself (Figures 3 and 4). The reason for this remains unclear, and is also despite increased immunogen doses being used for the third and fourth immunisations, but provides evidence that related toxins (batroxobin shares 93% and 62% amino acid percentage identity with ancrod and RVV-V, respectively) (43, 58, 59) may stimulate considerably different titres of specific antibodies. It was also noticeable, and perhaps unsurprising, that each of the experimental antivenoms generated against the recombinant toxins exhibited considerably reduced immunological binding levels to the crude venoms, and were outperformed by the ‘anti-venom mix’ control antivenom in this regard (Figure 3). These findings are not unexpected considering that snake venoms consist of a variety of distinct toxins, and thus our SVSP-focused antivenoms can only be expected to recognise a proportion (i.e. binding titres are likely to be somewhat proportional to SVSP content in each venom). This perhaps makes our findings observed with *C. rhodostoma* venom particularly noticeable, as the anti-ancrod, anti-RVV-V and the anti-toxin mix antivenoms produced end-point titration ELISA binding profiles nearly comparable to the anti-venom mix control antivenom against this venom (Figure 3).

It is promising that the immunological profiling described above also extended to the experimental antivenoms conferring protection against toxin functional activities. Each of the toxin-specific antivenoms were capable of inhibiting a pathologically relevant functional activity of the recombinant toxins (fibrinogenolysis), and also reduced the consumption of fibrinogen and the prolongation of the PT measured in plasma spiking experiments (Figures 5-7). Although these inhibitory effects were generally greater against the recombinant toxin immunogens than the crude venoms, again likely due to distinct non-SVSP toxins also contributing to these functional activities, reductions in fibrinogenolysis, fibrinogen depletion and PT prolongations were observed with each of the toxin-specific experimental antivenoms. To contextualise these findings, it is worth nothing that for the latter two experiments, the toxin-specific antivenoms outperformed the commercial antivenom controls. Collectively these findings strongly suggest that a single recombinant toxin is capable of stimulating desirable inhibitory antibody responses against snake venom toxins.

However, perhaps somewhat surprisingly, we failed to observe any discernible advantage in combining the three recombinant SVSP toxins together as immunogens. The ‘anti-toxin mix’ antivenom resulted in highly comparable binding titres and toxin specificities to the anti-ancrod and anti-RVV-V antivenoms against the various immunogens and the panel of native snake venoms used in this study (Figures 3 and 4 and Supplementary Figures 3 and 4), while this antivenom was inferior in inhibiting the activity of the toxins and venoms measured in the plasma-spiking functional assays (Figures 6 and 7). Thus, in this case, a single representative SVSP toxin was capable of generating at least comparable antibody responses to a mixture of multiple different SVSPs, which is an intriguing finding analogous to recent observations with crude venoms that fewer immunogens covering toxin diversity may be superior to more (27). However, for such an approach to be successful, the ‘right’ immunogen to select remains of critical importance, and relying on a single toxin to generate broad, yet toxin family-specific, antibody responses comes with considerable risk, and perhaps even more so given that use of recombinant batroxobin here resulted in an antivenom with poor immunological recognition of native venom toxins.

Despite the promise of the findings present herein, it is important to note that there are several limitations associated with this study. The most important of these is that, due to the restricted blood sample volumes collected post-immunsiation (which were due to the relatively small sample size per immunogen group (n=5) and ethical constraints), our analyses of the binding and functional neutralisation of SVSP toxins and crude venoms were limited to *in vitro* experiments. While there remain considerable issues associated with current preclinical models used to assess antivenom efficacy, including their questionable relevance for modelling VICC (60), these *in vivo* models remain the gold standard for efficacy testing. Thus, important next steps with recombinant toxin-specific antivenoms, such as those generated here, would be to test whether their functional inhibitory profiles are capable of conferring preclinical protection against venom-induced pathologies in appropriate *in vivo* models. In the context of the SVSP toxins targeted in this study, assessing whether experimental antivenoms designed against recombinant toxins can reduce systemic haemorrhage and/or the severity of coagulopathy would be valuable readouts.

The capability of toxin family-specific antivenoms to broadly protect against venom-induced lethality may be restrictive, particularly when attempting to neutralise snake venoms where distinct toxin families are contributing substantially towards severe pathology. However, a previous study that used a recombinantly expressed neurotoxin as immunogen, specifically a chimeric consensus short-chain *α* -neurotoxin three finger toxin, resulted in an equine experimental antivenom capable of neutralising the lethal effects of both specific neurotoxins and a variety of crude neurotoxic snake venoms (52). Thus, in future research it would be fascinating to explore whether recombinant ancrod or RVV-V, perhaps also along with an informatically designed consensus SVSP toxin, might be capable of generating an antivenom capable of providing a degree of preclinical protection against geographically diverse vipers that cause systemic haemotoxicity *in vivo*. Should such an approach provide limited efficacy, a strong case would likely remain for recombinant toxins retaining value for antivenom production in the form of their use as either supplemental immunogens to either ‘boost’ or redirect the immune response towards specific toxins of greatest pathological importance, or for generating separate polyclonal antibody pools that could be blended or used in conjunction with existing antivenom therapies in a fortifying approach. Overall, our findings further strengthen the potential value of using recombinant venom toxins as immunogens to stimulate focused and desirable antibody responses capable of neutralising specific venom-induced pathological effects. Such tools are a welcome addition to the diversity of experimental approaches currently being explored to circumvent limitations associated with current antivenoms, with the long-term goal of dramatically improving therapeutics for the world’s neglected tropical snakebite victims.

## Acknowledgments

The authors wish to thank Paul Rowley for expert snake husbandry at LSTM. N.A. acknowledges support from a PhD scholarship provided by the Ministry of Higher Education, Saudi Arabia and the Royal Embassy of Saudi Arabia, Cultural Bureau, London. N.R.C. acknowledges support from a Sir Henry Dale Fellowship (200517/Z/16/Z) jointly funded by the Wellcome Trust and Royal Society. This research was funded in part by the Wellcome Trust [200517/Z/16/Z]. For the purpose of open access, the author has applied a CC BY public copyright licence to any author accepted manuscript version arising from this submission.

