## Supplementary Information for "Exploring the utility of recombinantly expressed snake venom serine protease toxins as immunogens for generating experimental snakebite antivenoms"

**Supplementary Table 1. The snake venoms used in SDS-PAGE gel electrophoresis and immunoblotting experiments.**

| Species | Sub-family | Geographical region | Venom origin |
|---|---|---|---|
| ***Bitis arietans*** | Viperinae | sub-Saharan Africa | Nigeria |
| ***Bothrops asper*** | Crotalinae | Central America | Costa Rica |
| ***Bothrops atrox*** | Crotalinae | South America | Brazil |
| ***Bothrops jararaca*** | Crotalinae | South America | Brazil |
| ***Crotalus atrox*** | Crotalinae | North America | USA |
| ***Calloselasma rhodostoma*** | Crotalinae | Southeast Asia | Captive bred |
| ***Daboia russelii*** | Viperinae | South Asia | Sri Lanka |
| ***Deinagkistrodon acutus*** | Crotalinae | East Asia | Captive bred |
| ***Dispholidus typus*** | Colubrinae | sub-Saharan Africa | South Africa* |
| ***Echis carinatus*** | Viperinae | South Asia | India** |
| ***Echis ocellatus*** | Viperinae | West Africa | Nigeria |
| ***Rhabdophis subminiatus*** | Natricinae | Southeast Asia | Hong Kong |
| ***Trimeresurus albolabris*** | Crotalinae | Southeast Asia | Captive bred |

* Venom sourced commercially from Latoxan, France.

** Indian *Echis carinatus sochureki* venom (referred to as *E. carinatus* throughout) was collected from a single specimen that was inadvertently imported to the UK via a boat shipment of stone, and then rehoused at LSTM on the request of the UK Royal Society for the Prevention of Cruelty to Animals (RSPCA).

### (A) Nucleotide and amino acid sequences of ancrod

VIGGDECNINEHRFLVALYDSTTRNFLCGGVLIHPEWVITAKHCNKKSMVLYLGKHKQSVKFDDEQERFPKEKHFIRCNKPRTRWGEDIMLIRLNKPVNNSEHIAP

LSLPSNPPIVGSVCRVMGWGSINKYIDVLPDEPRCANINLYNYTVCRGVFPRIPKKSKILCAGDLQGRLDSCHCDSGGPLICSEEFHGIVYRGPNPCAQPDKPALYT

NIFDHLHWILSIMAGNATCYP

seq primer Fcmut1:5'

CCTAGACTCAGCCGGCTCTCCACGCTTTGCCTGACCCTGCTTGCTCAACTCTACGTCTTTGTTTCGTTTTCTGTTCTGCGCCGTTACAGATCCAAG

START

CTGTGACCGGCGCCTACCTGAGATCACCGGCGAAGGAGGGCCACCATGTACAGGATGCAACTCCTGTCTTGCATTGCACTAAGTCTTGCACTT

EcoRI

AATGCGAATTCGTCATTGGAGGTGATGAATGTAACATAAATGAACATCGTTTCCTTGTAGCCTTGTATGACAGTACGACTCGGAATTTTCTCTGTGGTGGG

GTTTTGATCCATCCGGAATGGGTGATCACTGCTAAACACTGCAACAAGAAAAGTATGGTCCTATACCTTGGTAAGCATAAACAAAGTGTAAAATTTGACGA

TGAGCAGGAAAGATTCCCAAAGGAGAAGCACTTTATTCGCTGTAACAAACCCCGTACCAGATGGGGCGAGGACATCATGTTGATCAGGCTGAACAAACCT

GTTAACAACAGTGAACACATCGCTCCTCTCAGCTTGCCTTCCAACCCTCCCATTGTGGGCTCAGTTTGCCGTGTTATGGGATGGGGCTCAATCAATAAATAT

ATAGACGTTTTGCCCGATGAACCTCGTTGTGCTAATATTAACCTGTACAATTACACGGTGTGTCGTGGAGTTTTTCCAAGGATACCAAAGAAAAGCAAAAT

ATTGTGTGCAGGTGACCTGCAAGGACGCCTAGATTCATGTCACTGTGACTCTGGGGGACCTCTCATTTGTAGTGAAGAATTTCATGGCATTGTATATCGGG

GACCCAATCCTTGTGCCCAACCAGATAAGCCTGCCCTCTACACCAACATCTTCGATCATCTTCACTGGATCCTTAGCATTATGGCAGGAAATGCAACTTGCT

ATCCACATCACCATCACCATCACTGAGCTAGCTGGCCAGACATGATAAGATACATTGATGAGTTTGGACAAACCA

6X his tag tail STOP/NheI seq primer Fcmut1:3'

### (B) Nucleotide and amino acid sequences of Batroxobin

MYRMQLLSCIALSLALVTEFVIGGDECDINEHPFLAFMYYSPRYFCGMTLINQEWVLTAAHCNRRFMRIHLGKHAGSVANYDEVVRYPKEKFICPNKKKNVITDK

DIMLIRLDRPVKNSEHIAPLSLPSNPPSVGSVCRIMGWGAITTSEDTYPDVPHCANINLFNNTVCREAYNGLPAKTLCAGVLQGGIDTCGGDSGGPLICNGQFQGI

LSWGSDPCAEPRKPAFYTKVFDYLPWIQSIIAGNKTATCP

seq primer Fcmut1:5'

CCTAGACTCAGCCGGCTCTCCACGCTTTGCCTGACCCTGCTTGCTCAACTCTACGTCTTTGTTTCGTTTTCTGTTCTGCGCCGTTACAGATCCAAG

START

CTGTGACCGGCGCCTACCTGAGATCACCGGCGAAGGAGGGCCACCATGTACAGGATGCAACTCCTGTCTTGCATTGCACTAAGTCTTGCACTT

EcoRI

AATGCGAATTCGTCATTGGAGGTGATGAATGTGACATAAATGAACATCCTTTCCTTGCATTCATGTACTACTCTCCCCGGTATTTCTGTGGTATGACTTTGAT

CAACCAGGAATGGGTGCTGACCGCTGCACACTGTAACAGGAGATTTATGCGCATACACCTTGGTAAACATGCCGGAAGTGTAGCAAATTATGATGAGGTG

GTAAGATACCCAAAGGAGAAGTTCATTTGTCCCAATAAGAAAAAAAATGTCATAACGGACAAGGACATTATGTTGATCAGGCTGGACAGACCTGTCAAAA

ACAGTGAACACATCGCGCCTCTCAGCTTGCCTTCCAACCCTCCCAGTGTGGGCTCAGTTTGCCGTATTATGGGATGGGGCGCAATCACAACTTCTGAAGAC

ACTTATCCCGATGTCCCTCATTGTGCTAACATTAACCTGTTCAATAATACGGTGTGTCGTGAAGCTTACAATGGGTTGCCGGCGAAAACATTGTGTGCAGG

TGTCCTGCAAGGAGGCATAGATACATGTGGGGGTGACTCTGGGGGACCCCTCATCTGTAATGGACAATTCCAGGGCATTTTATCTTGGGGAAGTGATCCC

TGTGCCGAACCGCGTAAGCCTGCCTTCTACACCAAGGTCTTTGATTATCTTCCCTGGATCCAGAGCATTATTGCAGGAAATAAAACTGCGACTTGCCCGCAT

CACCATCACCATCACTGAGCTAGCTGGCCAGACATGATAAGATACATTGATGAGTTTGGACAAACCA

6X his tag tail STOP/NheI seq primer Fcmut1:3'

### (C) Nucleotide and amino acid sequences of RVV-V

VVGGDECNINEHPFLVALYTSASSTIHCAGALINREWVLTAAHCDRRNIRIKLGMHSKNIRNEDEQIRVPRGKYFCLNTKFPNGLDKDIMLIRLRRPVTYSTHIAPV

SLPSRSRGVGSRCRIMGWGKISTTEDTYPDVPHCTNIFIVKHKWCEPLYPWVPADSRTLCAGILKGGRDTCKGDSGGPLICNGEMHGIVAGGSEPCGQHLKPAV

YTKVFDYNNWIQSIIAGNRTVTCPP

seq primer Fcmut1:5'

CCTAGACTCAGCCGGCTCTCCACGCTTTGCCTGACCCTGCTTGCTCAACTCTACGTCTTTGTTTCGTTTTCTGTTCTGCGCCGTTACAGATCCAAG

START

CTGTGACCGGCGCCTACCTGAGATCACCGGCGAAGGAGGGCCACCATGTACAGGATGCAACTCCTGTCTTGCATTGCACTAAGTCTTGCACTT

EcoRI

AATGCGAATTCGTCGTTGGAGGTGATGAATGTAACATAAATGAACATCCTTTCCTTGTAGCCTTGTATACCTCTGCCTCTAGCACGATTCACTGTGCTGGTG

CTTTGATCAACAGGGAATGGGTGCTCACCGCTGCACACTGTGACAGGAGAAATATCCGGATAAAGCTTGGTATGCATAGCAAAAATATACGAAATGAGG

ATGAGCAGATAAGAGTCCCAAGGGGCAAGTACTTTTGTCTTAATACCAAATTCCCCAACGGATTAGATAAGGACATCATGTTGATCAGGCTGAGAAGACC

TGTTACCTACAGTACACACATCGCGCCTGTCAGCTTGCCTTCCCGTTCTCGCGGTGTGGGCTCACGTTGCCGTATTATGGGATGGGGCAAAATCTCAACTAC

TGAAGATACTTATCCTGATGTCCCTCATTGTACTAACATCTTCATAGTCAAGCATAAGTGGTGTGAACCACTTTATCCATGGGTGCCTGCTGATAGCAGAAC

ATTGTGTGCTGGTATCCTAAAAGGAGGCAGAGATACATGTAAGGGTGACTCTGGGGGACCGCTCATCTGTAATGGAGAAATGCACGGCATTGTAGCTGG

GGGGTCTGAACCTTGTGGCCAACATCTTAAACCTGCTGTCTACACCAAGGTCTTCGATTATAATAACTGGATCCAGAGCATTATTGCAGGAAATAGAACTG

TGACTTGCCCCCCGCATCACCATCACCATCACTGAGCTAGCTGGCCAGACATGATAAGATACATTGATGAGTTTGGACAAACCA

6X his tag tail STOP/NheI seq primer Fcmut1:3’

**Supplementary Figure 1. Selection of SVSPs toxins for recombinant expression.** Three SVSPs toxins were selected for recombinant expression as biologically active and functionally relevant components of distinct viper venoms. Coding sequences were sourced from the GenBank database of the National Centre for Biotechnology Information and signal peptides were removed. **(A)** Ancrod from *C. rhodostoma* (GenBank: L07308.1), **(B)** Batroxobin from *B. atrox* (GenBank: J02684.1) and **(C)** RVV-V from *D. russelii* (GenBank: MF289120.1). The restriction enzymes used were EcoRI (cut site: 5’-GAATTC-3’) and NheI (cut site: 5’-GCTAGC-3’). 6X his tag tail (CATCACCATCACCATCAC). Primers used were Fcmut1:5’ (ACCCTGCTTGCTCAACTCT) and Fcmut1:3’ (TGAGTTTGGACAAACCA).

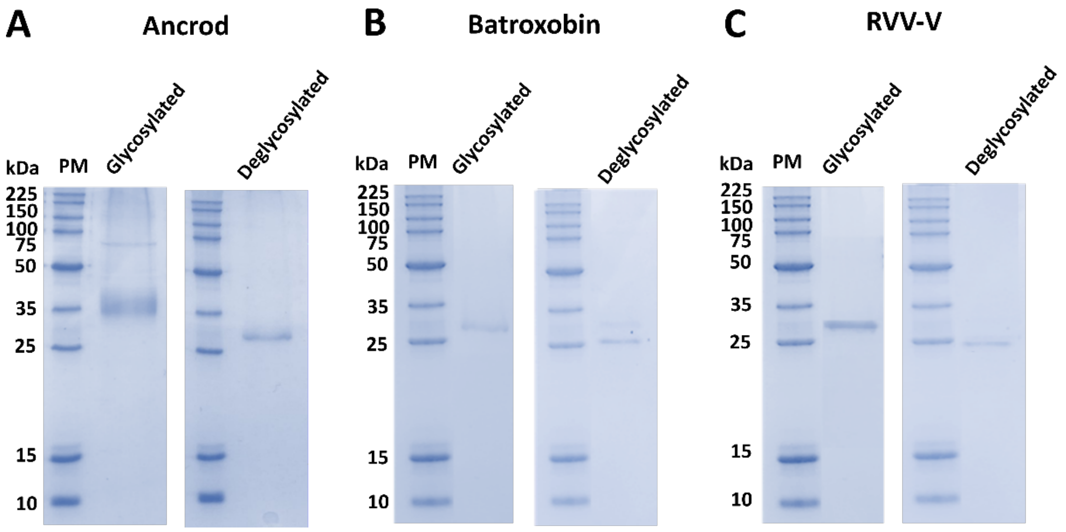


**Supplementary Figure 2. Coomassie-stained SDS-PAGE gels of glycosylated/deglycosylated recombinant toxins.** Glycosylated/deglycosylated recombinant toxins (all 7 µg) were separated by reduced 15% SDS-PAGE gel electrophoresis and visualised by Coomassie blue staining. Deglycosylated recombinant toxins (7 µg) were generated via incubation with PNGase F at 37 °C overnight. **(A)** Ancrod, **(B)** Batroxobin, and **(C)** RVV-V. For each, the following layout was used: Lane 1, protein marker (PM); Lane 2, glycosylated recombinant toxin; Lane 3, protein marker; Lane 4, deglycosylated recombinant toxin.

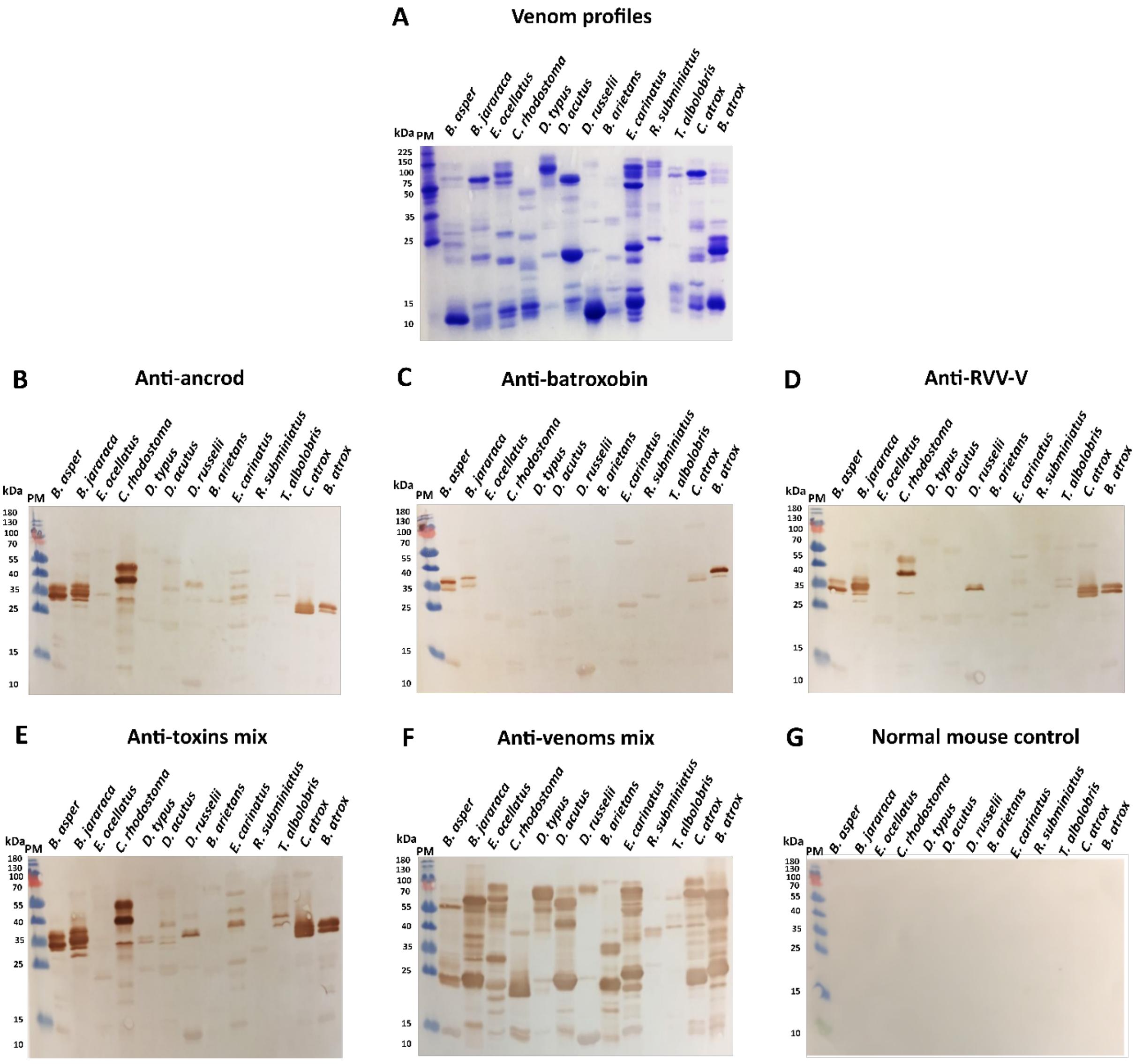


**Supplementary Figure 3. Immunological recognition of proteins found in a diverse array of haemotoxic snake venoms by the experimental antivenoms. (A)** Snake venoms were separated by reduced 15% SDS-PAGE gel electrophoresis and visualised by Coomassie blue staining. The same venom samples were transferred to nitrocellulose membranes for immunoblotting experiments and incubated with 1:5,000 dilutions of primary antibodies of each of the experimental antivenoms, specifically: **(B)** anti-ancrod, **(C)** anti-batroxobin, **(D)** anti-RVV-V, **(E)** anti-toxins mix, **(F)** anti-venoms mix (as positive control) and **(G)** normal mouse control (as negative control). PM indicates protein marker.

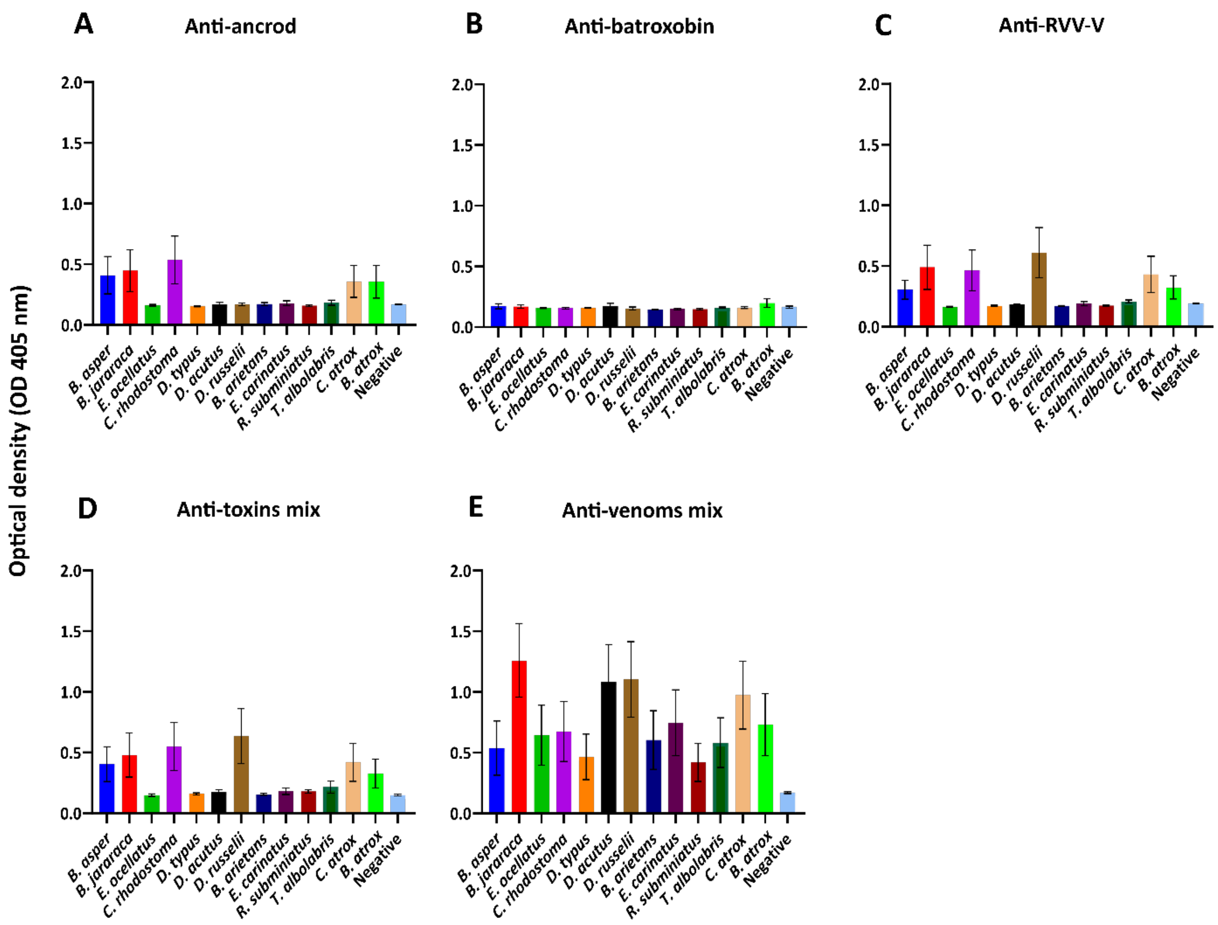


**Supplementary Figure 4. Immunological cross-reactivity of the experimental antivenoms with a diverse array of haemotoxic snake venoms.** The various experimental antivenoms were standardized to a 1:5,000 dilution and used as primary antibodies with the mean data shown and error bars representing standard deviation (SD) of the duplicate measurements. Data shown represents binding levels by: **(A)** anti-ancrod, **(B)** anti-batroxobin, **(C)** anti-RVV-V, **(D)** anti-toxins mix and **(E)** anti-venoms mix (as positive control) experimental antivenoms. Normal mouse control was used as a negative control and is displayed in each panel.

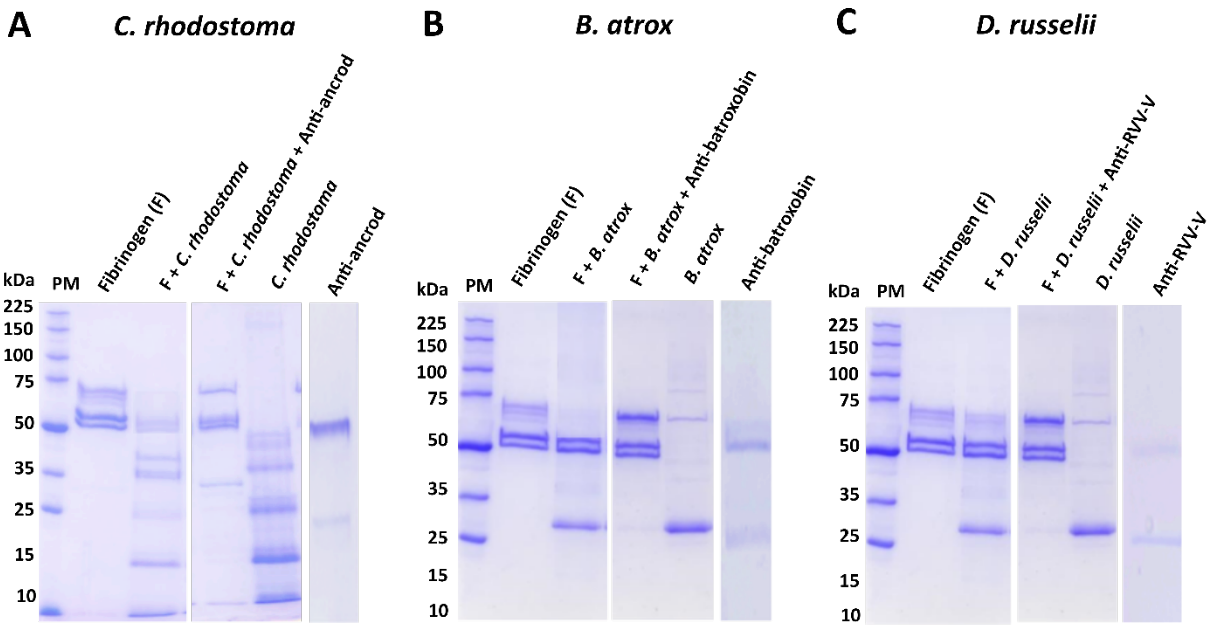


**Supplementary Figure 5. Experimental antivenoms directed against recombinant SVSP toxins show a degree of inhibition against the fibrinogenolytic activity of snake venoms.** Degradation SDS-PAGE gel electrophoretic profiles are displayed following the incubation of various samples at 37 °C for 120 min. Panels show different data obtained with the different snake venoms used as controls: **(A)** *C. rhodostoma*, **(B)** *B. atrox,* and **(C)** *D. russelii*. For each, the following layout was used: Lane 1, protein marker (PM); Lane 2, human fibrinogen (3 µg); Lane 3, fibrinogen + venom (7 µg, *C. rhodostoma*, *B. atrox* or *D. russelii*); Lane 4, fibrinogen + venom + experimental antivenom (0.5 µg, anti-ancrod, anti-batroxobin or anti-RVV-V); Lane 5, venom only; Lane 6, specific experimental antivenom only.

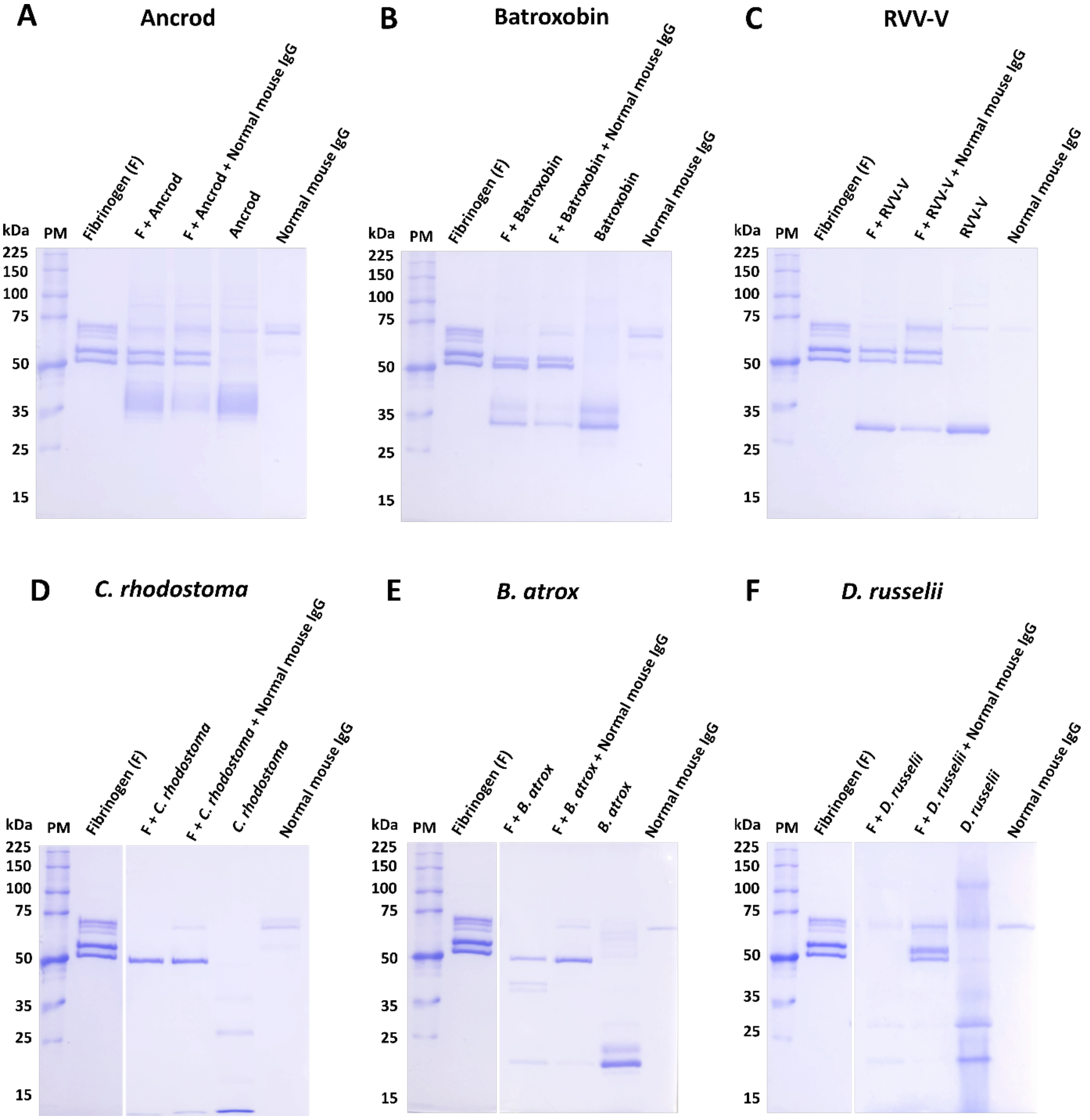


**Supplementary Figure 6. Normal non-immunised mouse IgG provides no protective effect against the fibrinogenolytic activity of recombinant toxins and snake venoms.** Degradation SDS-PAGE gel electrophoretic profiles are displayed following the incubation of various samples at 37 °C for 120 min. Panels show different data obtained with the different recombinant toxins and snake venoms used as immunogens: **(A)** ancrod, **(B)** batroxobin, **(C)** RVV-V, **(D)** *C. rhodostoma* venom, **(E)** *B. atrox* venom, and **(F)** *D. russelii* venom. For each, the following layout was used: Lane 1, protein marker (PM); Lane 2, human fibrinogen (3 µg); Lane 3, fibrinogen + toxin/venom (7 µg, ancrod, batroxobin, RVV-V, *C. rhodostoma*, *B. atrox* or *D. russelii*); Lane 4, fibrinogen + toxin/venom + normal mouse control (0.5 µg, 1 mg/ml); Lane 5, toxin/venom only; Lane 6, normal mouse control IgG only.

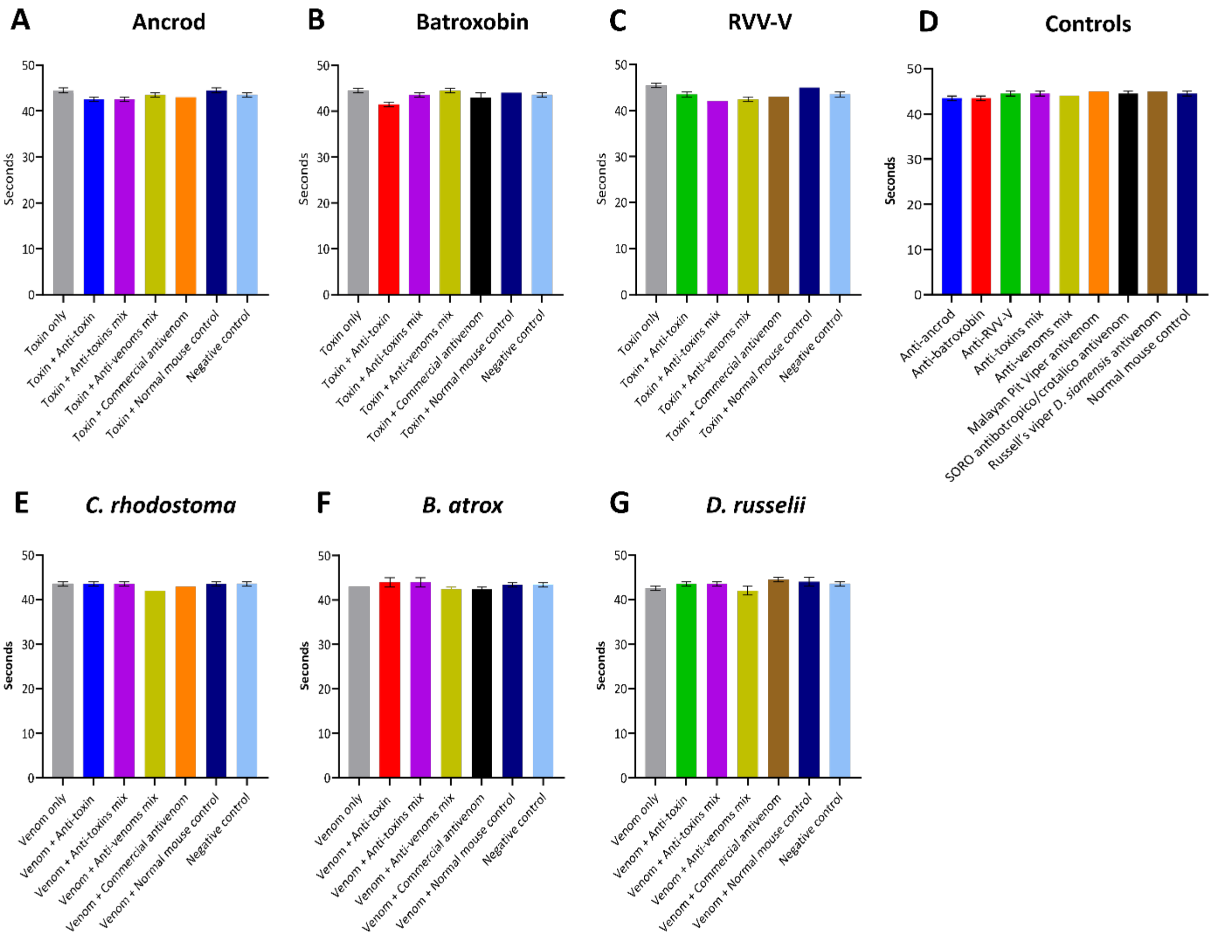


**Supplementary Figure 7. Inhibition of coagulation disturbances defined by the activated partial thromboplastin time (aPTT).** The assay measured the combined effect of the clotting factors of the intrinsic and common coagulation pathways (in seconds) in the presence of the recombinant toxins and their recovery effect by adding specific experimental antivenoms/specific commercial antivenoms (Monovalent equine antivenom for Malaysian Pit Viper against ancrod and *C. rhodostoma*, SORO antibotropico/crotalico antivenom against batroxobin and *B. atrox* and monovalent equine *D. siamensis* commercial antivenom against RVV-V and *D. russelii*) and the normal mouse control incubated with FFP. Data shown represents the following immunogens: **(A)** Ancrod, **(B)** Batroxobin, **(C)** RVV-V, **(D)** the controls**, (E)** *C. rhodostoma* venom, **(F)** *B. atrox* venom and **(G)** *D. russelii* venom. Each experimental antivenom alone, each commercial antivenom alone and normal mouse control alone were used as negative controls. Error bars represent the standard deviation (SD) of duplicate measurements.
